# Improved marker detection for rare population in single-cell transcriptomics through text mining-inspired scoring approach

**DOI:** 10.1101/2025.11.10.687759

**Authors:** Jinjin Chen, Mengbo Li, Dharmesh D. Bhuva, Melissa J. Davis, Anthony T. Papenfuss, Chin Wee Tan

## Abstract

Accurate identification of cell types and states is a crucial step when analysing single cell RNA-seq data. Inaccurate cell type annotation leads to spurious results and biological interpretations in subsequent analyses. Cell type identification is traditionally done using established marker genes of each cell population. However, many existing methods do not perform well for imbalanced cell populations, which often occurs in the presence of rare cell types. Most existing methods tend to be biased towards the dominant cell populations or highly expressed genes, leading to inaccurate results. Here, we introduce the package *smartid*, which accurately identifies markers from even imbalanced data across batches by employing a modified Term Frequency-Inverse Document Frequency approach with Gaussian Mixture Model. *smartid* is also a gene-set scoring method which is able to distinguish the target group of interest. *smartid* is implemented in R and is freely available on Bioconductor at https://bioconductor.org/packages/release/bioc/html/smartid.html.

## Main

Single cell RNA-seq (scRNA-seq) has improved our understanding of cellular heterogeneity, revealing intricate patterns of gene expression for individual cells. Delving deeper into the molecular structure of tissues and organs as well as the interactions and signalling pathways between different cell types, marker gene identification has become critical and has gained significant attention in recent years (1–3). Marker genes are widely used across biological and medical research. Foremost, accurate identification of cell type marker genes can help annotate cells into different biologically meaningful types or subpopulations (4). In the context of disease, comparing healthy and diseased samples enables the identification of prognostic markers, which can then be used as diagnostic indicators (5). Potential therapeutic targets might be found by identifying the markers specific to cancer cell populations, particularly those responsive to specific drugs (6).

One challenge to accurately identifying marker genes is dealing with imbalanced data, where certain populations or genes are overrepresented, leading to biased results towards these dominant groups or genes (7,8). Such imbalanced data, characterized by varying cell numbers across different cell types, poses significant difficulties for marker gene identification. First, it reduces the statistical power to detect the genes associated with the underrepresented or rare populations. Algorithms used to identify marker genes may not be able to adequately account for the variability present in smaller populations, thus weakening their analytical power. Second, imbalanced data can lead to overfitting of machine learning classifiers, where the model becomes biased towards the dominant populations due to their higher representation in the dataset. This results in the model performing well on these overrepresented classes but poorly on less frequent ones, ultimately reducing its overall predictive accuracy and generalizability. Most current methods do not adequately address this issue, often focusing on the dominant cell populations and highly expressed genes while unable to account for rare cell populations or lowly expressed yet highly variable genes. To overcome this challenge, we developed an R/Bioconductor package *smartid*, which takes into account all cell populations and genes in the data.

This work describes the software package *smartid* (Scoring and MARker identification approach based on TF-IDF), a novel approach for gene-set scoring and marker gene identification on single-cell data. Term frequency-inverse document frequency (TF-IDF) is a popular method in natural language processing, originally developed to quantify the importance of terms in a document (9). It consists of two factors: (1) term frequency (i.e., TF), which measures the frequency of each word in a document, and (2) inverse document frequency (i.e., IDF), which is used to down-weight the terms that appear often in the corpus and to assign a higher weight to those words that are rare across the entire corpus but appear in the particular documents. Applying this analogy onto transcriptomic data (genes as words and cells as documents), TF-IDF can be used to identify and highlight the genes that are highly relevant for the classification. It has been successfully applied to many domains, including text mining and information retrieval, proving its utility for feature selection (9–11). In this study, we developed *smartid* to identify marker genes in scRNA-seq data, especially for rare cell populations.

Here, we demonstrate that *smartid* achieved good performance on both simulated and real-life biological data. When benchmarked against other commonly used methods, we showed *smartid* exhibits improved performance compared to other methods as a marker gene identification method. *Smartid* is also able to discriminate closely similar cell types when scoring cells based on the specific marker genes. Beyond scRNA-seq applications, our analysis of image-based spatial transcriptomics datasets revealed that *smartid* markers can accurately capture tissue structures in the data as seen from ground truth biological samples. The spatial patterns we observed matched what was expected in biological tissue architecture, which increases confidence in the markers’ biological validity. This suggests that *smartid* also holds promising applications in the rapidly evolving spatial omics, making it a useful tool for single-cell data analysis in multiple fields. Overall, *smartid* is a user-friendly, fast and accurate marker gene identification approach in the field of bioinformatics, particularly helpful in providing insights into rare cell populations.

## Results

### Overview of *smartid* framework

The *smartid* framework represents a concerted approach for automated identification of signature markers in complex biological datasets, with particular efficacy for detecting markers in rare cell populations from imbalanced data. The *smartid* framework integrates several core algorithmic components to enable robust signature detection and scoring from high-dimensional omics data. *smartid’s* algorithm retains the core principles of TF-IDF while incorporating specific mechanisms to mitigate the impact of highly sparse data on gene weights. Gene weight is a metric used to quantify the importance of a gene in the data for a particular classification or prediction result (e.g. cell type and drug response), which is very important during feature selection. The whole workflow of *smartid* is divided into 4 steps: scoring, scaling, transforming and selecting (**Figure 1**). Depending on whether the input data contains the cell/sample label, either labelled scoring or unlabelled scoring approach should be performed. The labelled score will be scaled to avoid bias towards large numerical values. The scaled score will then be transformed into an importance for each gene within each cell group using a softmax function, with a range from 0 to 1. Finally, the marker gene selection of *smartid* is done by modelling the mixtures of parametric distributions using the expectation-maximization (EM) algorithm (12).

**Figure 1.**
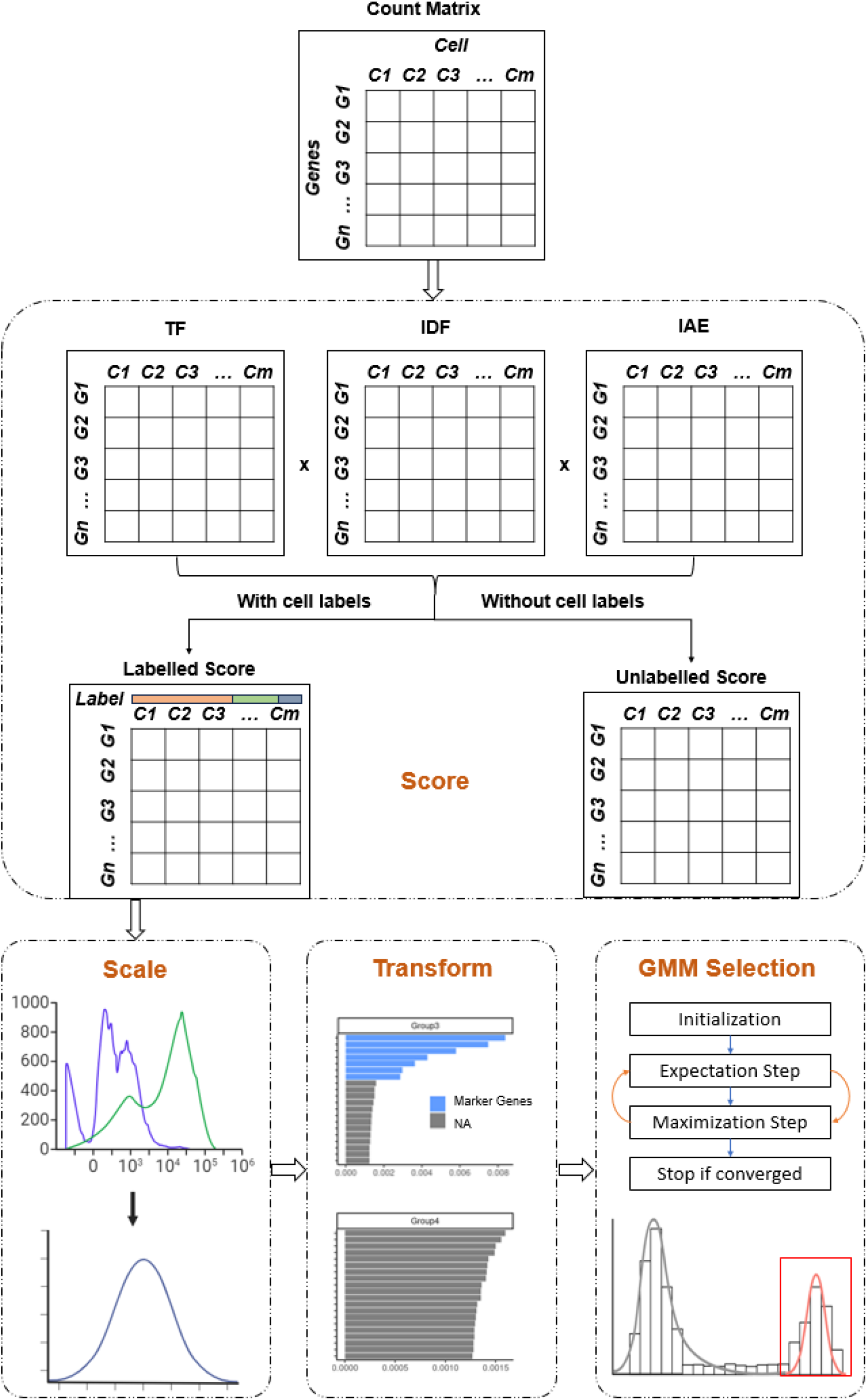
*smartid* schematic workflow. The workflow for *smartid*: (1) scoring, conduct labelled or unlabelled scoring based on data availability, (2) scaling, use a weighted mean to scale scores to reduce bias from large numerical values, (3) transforming, use softmax function to transform scores into importance, (4) selecting, using a Gaussian Mixture Model (GMM) with a default threshold of 0.99 to identify marker genes for each cell group.

Essentially, *smartid* leverages a modified TF-IDF specifically adapted for single-cell data for feature weighting, enhancing the distinction between biologically relevant signals and noise. Unlike conventional differential expression methods, this approach enables the detection of signature markers that show distinctive expression patterns across biological groups, even when these markers don’t exhibit high expression levels. By adding a new factor IAE (Inverse Average Expression), *smartid* is able to identify the markers that are widely expressed in general but highly expressed in a specific population. Different modifications were made to TF-IDF-IAE to properly accommodate the labelled or unlabelled input data. (see details in **Methods**)

Scaling is necessary before the feature selection as different features typically share differing ranges of expression. However, scaling imbalanced data is a big challenge. Consider a dataset containing both a rare class and a dominant class. When the traditional z-score scaling method is applied, the characteristics of the dominant class, such as its mean and variance, will disproportionately affect the scaling process. Due to this bias, the scaled score within the dominant class tends to cluster around zero. As a result, genes that are highly expressed in the dominant class may show unexpectedly low z-scores, even if they are biologically significant within that class. To mitigate this problem caused by the imbalanced data, we performed a weighted scaling to make each class contribute equally at the class level. A non-negative importance of each gene to each class will then be computed after softmax transformation. (Details in **Methods**)

To improve the detection, *smartid* incorporates Gaussian Mixture Model (GMM) approach for clustering, enabling unsupervised classification of heterogeneous biological markers within complex expression data distributions. This probabilistic framework accommodates the inherent variability in gene expression data, resulting in more robust signature identification even in heterogeneous cellular environments.

Technical variations and batch effects are usually inherent present in large-scale biological experiments. These effects that’s present in multi-sample datasets can introduce significant variability in cellular expression profiles within identical groups or populations, potentially compromising the fidelity of marker identification. This variability poses a substantial challenge to accurate marker identification. To mathematically account for technical artifacts while preserving biological signals, *smartid* incorporates a generalized linear model (GLM) approach using design matrices incorporating coefficients for experimental conditions and batch identifiers.

This integrated approach results in a comprehensive tool for analysing complex large-scale datasets, particularly valuable for investigating cellular heterogeneity, developmental trajectories, and disease-specific molecular signatures in contexts where rare cell populations or subtle expression changes are of primary interest.

### *smartid* identifies highly group-specific marker genes on simulated data

To evaluate the performance of *smartid*, it was applied to a scRNA-seq data simulated by *splatter* (13), showing its high accuracy in the identification of real marker genes. The data was simulated as described in **Data and methods** with 1,000 genes and 3,000 cells divided into 4 groups. *smartid* successfully identified the correct marker genes (**Figure 2B**) as validated by the ground truth differential expressed genes (DEGs) (**Figure 2A**). The coloured true DEGs (not grey) are prominently ranked at the top of each corresponding group by *smartid* importance, and there is a significant difference in the distribution of the importance between the groups with (Groups 1 to 3) and without DEGs (Group 4) (**Figure 2A** and **B**). Looking at Groups 1 to 3, a clear transition between DEGs (coloured) and non-DEGs (grey) was observed to effectively separate them (in **Figure 2A**). In contrast, Group 4, having no DEGs, showed relatively smooth changes in importance across the genes, with no obvious mixtures in the distribution. This characteristic can be utilized to help determine the presence of specific marker genes within the group. The marker genes identified by *smartid* (**Figure 2B**) presented similar results to real DEGs (in **Figure 2A**), showing a clear separation between identified marker genes and non-marker genes. It is clear that the *smartid*-identified marker genes closely matched the ground truth (DEGs in each group), with no marker genes identified for Group 4 (**Figure 2C**).

**Figure 2.**
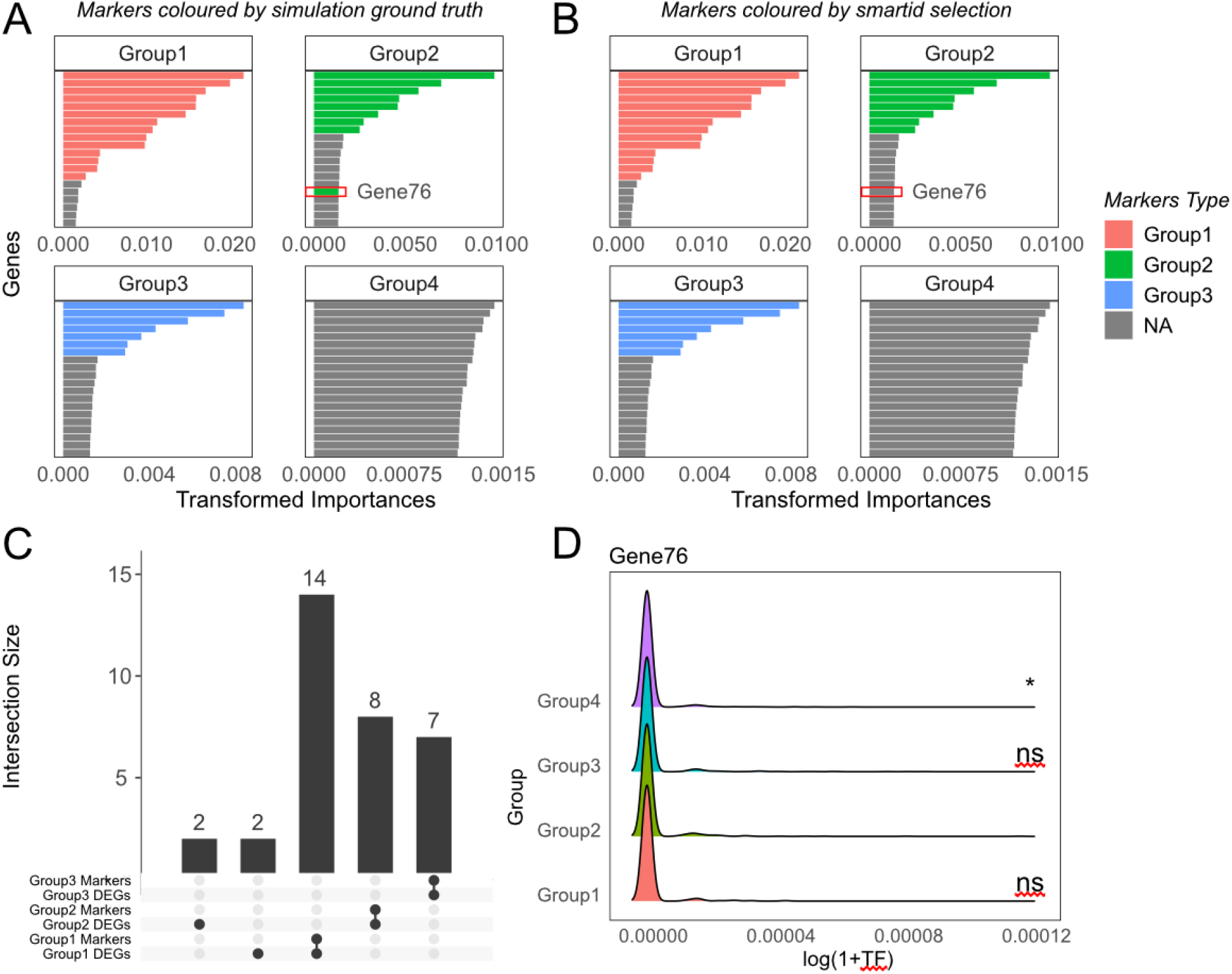
Performance of *smartid* on one simulated scRNA-seq data. (A)&(B) Bar plot of genes ranked by *smartid* transformed importance within each cell group. Genes are coloured by true DEGs of each group in (A) and by *smartid*-identified marker genes of each group in (B). (C) Overlap between *smartid*-identified marker genes and true DEGs for Group 1-3 (Group 4 has no DEGs). (D) Density plot of the log(1+TF) of ‘Gene76’ in all cells across groups. T statistical test was performed with reference to Group 2, ‘ns’ for not significant, ‘*’ for p < 0.05.

However, there was one particular DEG in Group 2, ‘Gene76’, that was not selected by *smartid* (**Figure 2A** and **B**). Based on the expression of this gene between groups (in **Figure 2D**), although it was named as one of the DEGs in Group 2, its actual expression level was very low with no clear transition in differences between different groups, thus the reason it failed to be identified by *smarted*, and rightfully so. The remaining DEGs not identified by *smartid* (in **Figure 2C**) showed similar results. This further validated the specificity and reliability of the markers identified by *smartid*.

### *smartid* offers improved performance in marker identification on simulated data

To assess the performance of *smartid* in marker gene identification, we performed a benchmark study against commonly used methods in the field including *Seurat* (14), *Monocle3* (7), *scran* (15) and *CelliD* (16) on a scRNA-seq data simulated by *splatter* (13), using testing parameters of 3,000 cells with 10,000 genes. The performance was evaluated by computing the F1 score, precision ratio, false discoveries in the negative control (Group 4) and the computational running time with the simulation repeated 1,000 times for robustness. Notably for *Monocle3*, 413 simulations failed to converge when applying *Monocle3* using default settings, thus only the results of the remaining 587 simulations were used. In this benchmark (**Supp Figure 2**), *smartid* and *scran* both appears to perform equally well and are generally significantly better than the other methods across all metrics. Both methods exhibited high accuracy and low false-positive discoveries, while possessing high computational efficiency on a dataset of 3,000 cells and 10,000 genes. Furthermore, *smartid* and *scran* maintain a stable performance across diverse groups regardless of population size. In comparison, *CelliD, Monocle3* and *Seurat* demonstrate performance that correlates with group size, showing improved metrics with increasing group sizes (from Groups 1 to 3). This result suggests that population size may be a critical factor influencing the performance of these widely used methods when applying to imbalanced datasets.

To further assess whether the performance of these methods can be upheld on imbalanced data, we reran the simulation 1,000 times, with Group 1 containing only 1% of cells, while Groups 2 to 4 were each 20%, 30%, and 49%. Once again for *Monocle3*, 378 simulations failed and were removed. The results of the new benchmark again show that the performance of all methods decreased in Group 1 when there was a rare population in the data, but in Groups 2 and 3, *smartid* showed the best performance (**Supp Figure 3A**). Notably, on the 10k-cell dataset, *smartid* is comparable to *Seurat* in terms of computational efficiency, and significantly outperformed *scran* (**Supp Figure 3**).

Overall, the results (**Table 1**) suggest that *smartid* showed overall higher precision and F1 score than the other methods on imbalanced data, particularly for Groups 2 and 3, whose F1 scores were the only ones greater than 0.8. The values for precision were over 0.85 and the false discoveries were 0, indicative that the marker genes identified by *smartid* are highly specific for the target cell populations, suggesting that *smartid* is effective in distinguishing between marker and non-marker genes, with very few false positives. *CelliD, Monocle3* and *Seurat* all reported a significant number of false positives in the negative control Group 4, where no DEGs were expected to be present. By contrast, *smartid* is the only method that consistently outputs no false positives with an overall highest F1 scores and low standard deviations (SD) across groups, suggesting reduced sensitivity to dataset imbalance and improved ability to minimize false discoveries, even in the absence of true differential expression signals.

**Table 1.**
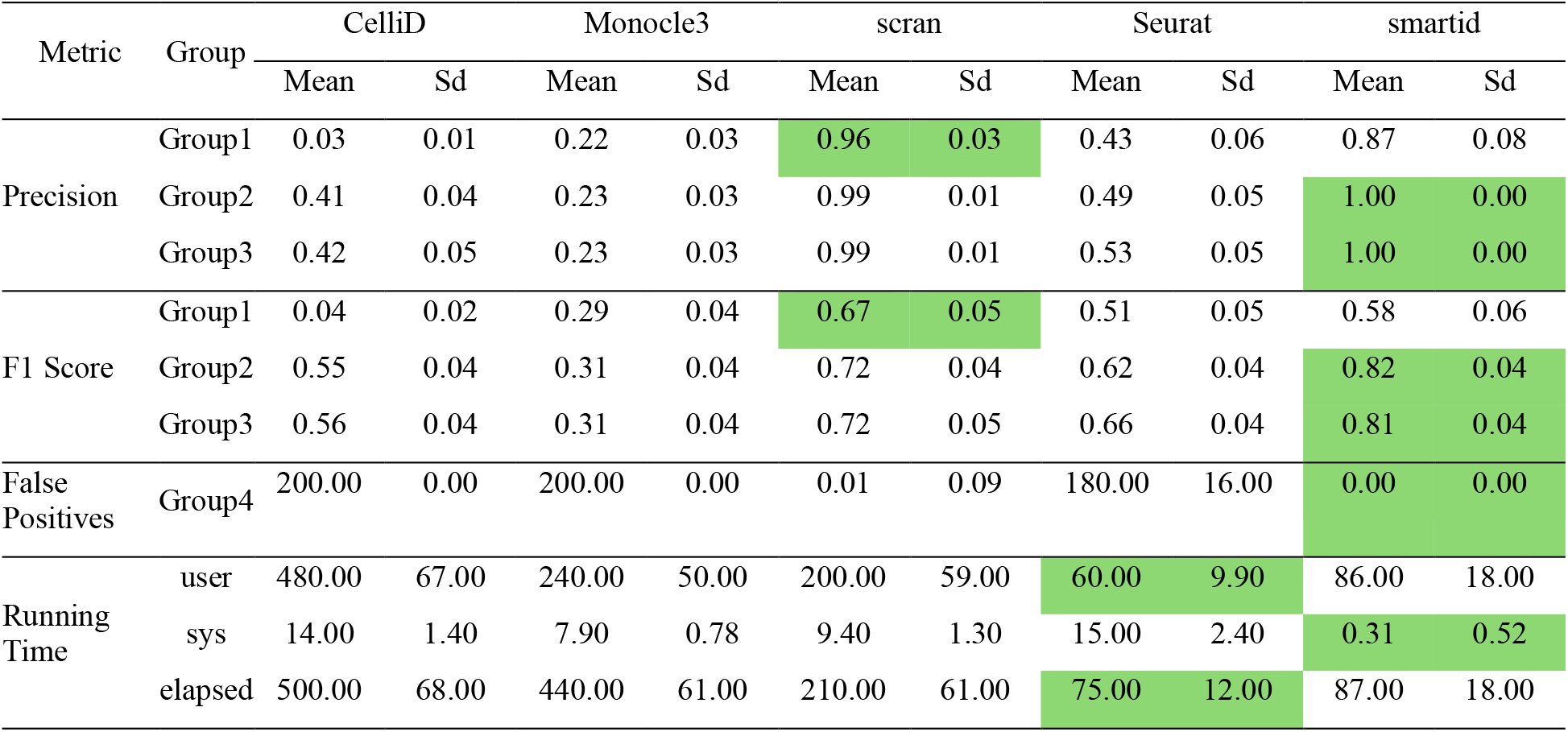
Performance of methods on simulated scRNA-seq data (10k cells).

*scran* exhibits similar performance to *smartid* in general, which might be assisted by the fact that the package *splatter* used to simulate the scRNA-seq data, assumes that the data strictly follows a negative binomial distribution, which aligns with the main algorithmic hypothesis of *scran*.

However, in terms of computational efficiency, *smartid* only required a slightly longer processing time compared to *Seurat* but is significantly faster than all other methods (*scran* requires at least twice the running time as *smarted*). As such, *smartid* provides a favourable balanced approach amongst the methods evaluated in this study. Detailed performance comparisons and computational times are shown in **Supp Figure 3**.

### *smartid* offers improved performance in marker identification on real data

To better understand the application and performance of these methods on real-life biological data, we applied them on a well-annotated dataset, ‘pbmc3k.final’ (17), and validated the superior performance of *smartid* over other methods. In this analysis, Monocle3 was not included due to its long running time and inadequate performance and *Seurat* appears to have identified the largest number of unique marker genes (compared to other methods, **Supp Figure 4**). However, previously eluted in the benchmarking results, these may include a large number of false positives and coupled with the poor ability of *Seurat* in distinguishing the target cell type (in **Supp Figure 5**), particularly between the similar (**Supp Figure 5C)** and rare cell types (**Supp Figure 5G)**. In contrast, this is not the case for *smartid* which performed consistently well across all cell types, particularly in discerning between CD8 T cells and NK cells **(Supp Figure 5C & G**). Importantly, *scran*, which performed well on the simulated data, did not perform well and appears to perform worse than *CelliD* in this real dataset, especially in the identification of the CD8 T cell marker (**Supp Figure 5C)**.

To further evaluate these four methods quantitatively, we computed the AUPRC based on the marker gene scores by different methods (**Figure 3A)**. *smartid* had the highest average AUPRC and the lowest SD with *scran* having the worst average AUPRC for marker gene identification. This indicates both the superior performance and reliability of *smartid*.

**Figure 3.**
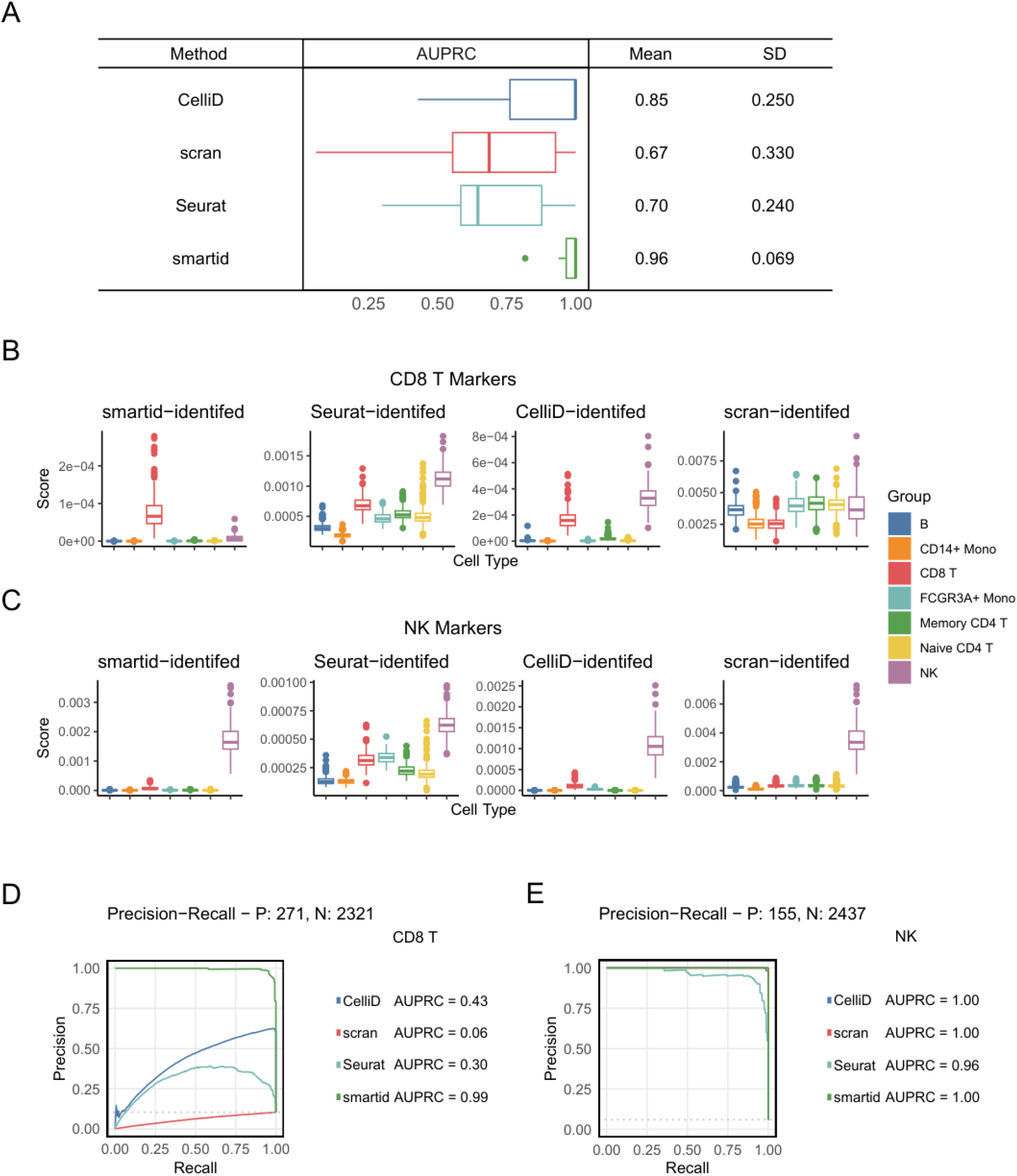
Performance of different methods on pbmc3k.final. (A) Summary of AUPRC based on *smartid* score, scores were calculated using marker genes identified by different methods for each cell type, AUPRC was calculated by comparing the *smartid* score to the binarized label for each cell type (e.g. NK cell and non-NK cell). (B)-(C) Boxplot of *smartid* score using CD8+ T cell marker genes (B), and NK cell marker genes (C), identified by different methods (left to right: *smartid, Seurat, CelliD* and *scran*). Boxes are coloured by cell type. (D)-(E) Precision-recall curve comparing the performance of the *smartid* score against the binarized cell labels for CD8+ T cells (D) and NK cells (E). The x-axis represents the recall, while the y-axis denotes the precision. Curves are coloured by the methods used to produce the marker genes.

Focusing on the performance at the cell type level, *smartid* showed the best results, especially when distinguishing between similar and rare cell types. In the presence of similar cell types, CD8 T and NK cells (the smallest population in ‘pbmc3k.final’), *smartid* still effectively identified marker genes highly specific to CD8 T cells and NK cells respectively, with a high AUPRC (> 0.9) (**Figure 3B-E**). In comparison, other methods yielded elevated scores for NK cells compared to CD8 T cells with a low AUPRC (< 0.5) for CD8 T cell markers (**Figure 3B&D**), suggesting difficulties in distinguishing between these two similar cell types. These results highlight the advantage of *smartid* over the existing methods on an existing biological dataset, reflected in both the ability to discrimination between similar cell types and the accurate marker gene identification for rare cell types.

### *smartid* improves the scoring-based group discrimination

To evaluate the performance of *smartid* as a gene-set scoring approach, we compared it (with or without labels) with four commonly used scoring methods: *UCell* (18), *GSVA* (19), *ssGSEA* (20) and *AUCell* (21), using the ‘pbmc3k.final’ dataset. All scoring methods achieved their highest scores for the corresponding cell types (**Figure 4A-F**, diagonals) using the same marker genes, indicating a clear identification of the target cell types.

**Figure 4.**
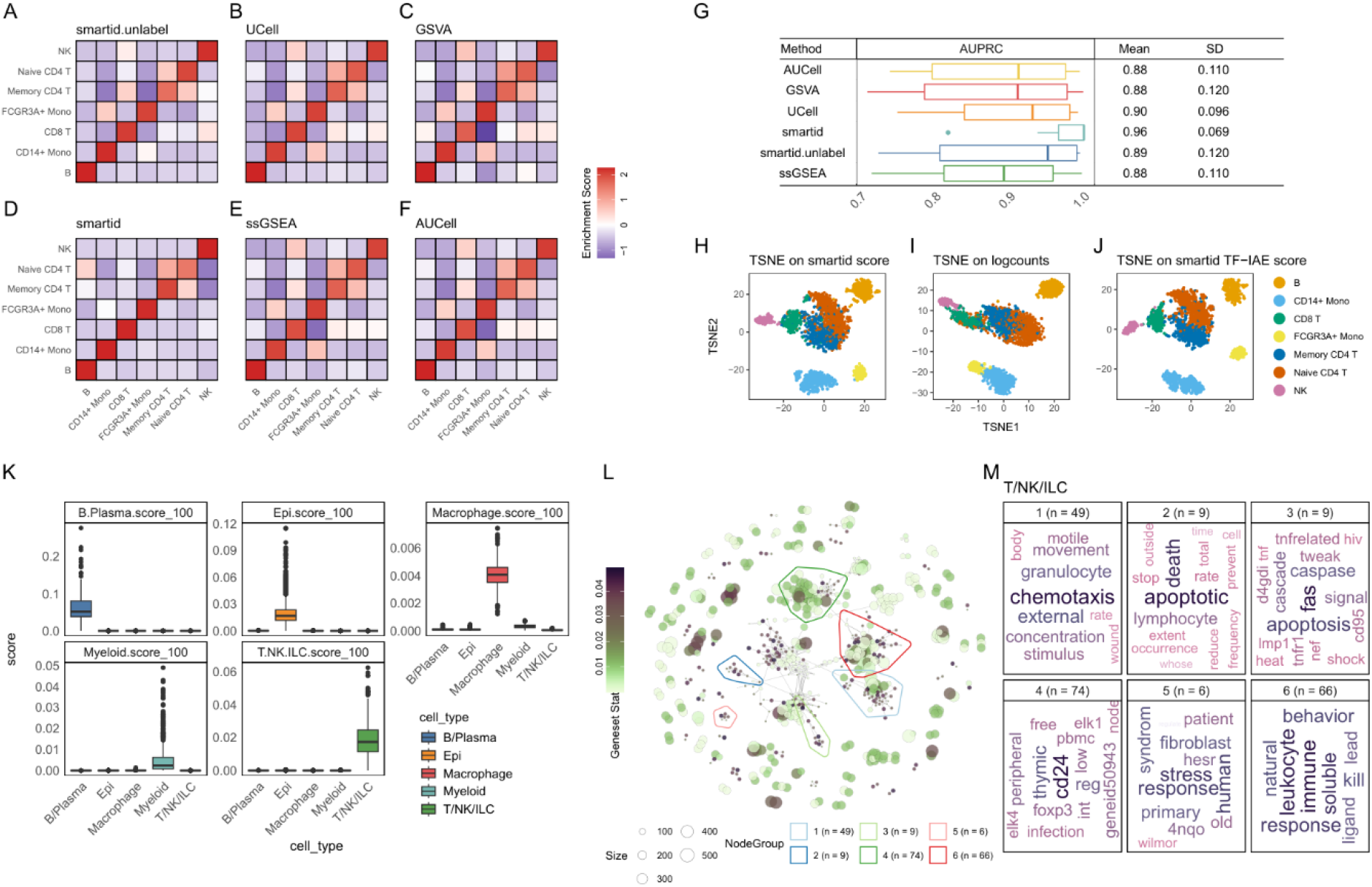
Performance of different scoring methods on pbmc3k.final and GSEA network visualization on scCOVID19 using *smartid*-derived marker genes. (A)-(F) Heatmap of gene-set scores (y axis in row) across different cell types (x axis in column) using different scoring methods. Each cell in the heatmap represents the score for a particular marker list in a particular cell type, calculated using different scoring methods with the marker genes identified by *smartid*. All the scores are scaled by row, the colour gradient indicates the magnitude of the score, from higher (red) to lower (blue). (G) Summary of AUPRC based on the scores computed by different methods using the same *smartid*-identified marker genes for each cell type. As described previously, AUPRC was calculated by comparing the score with the binarized cell type label for each cell type. (H)-(J) tSNE plot of pbmc3k.final dataset based on *smartid* TF-IDF-IAE score (H), *scran*-normalized log-counts (I) and *smartid* TF-IAE score (J). (K) Boxplot of *smartid* score across cell types using the cell type markers identified by *smarted*. Boxes are coloured by cell type. (L) Network visualization of the significantly enriched gene-sets based on T-NK-ILC cell marker genes, the top 6 clusters of the gene-sets (clustered by biological similarity) are highlighted with coloured circles. (M) Word clouds of the gene-set names from the top 6 clusters in (L). The number n in the bracket is the number of gene-sets in each cluster.

However, apart from the two CD4 T cell subtypes (naïve CD4 T cells and memory CD4 T cells), where the scores were consistently close between the two across all methods, the other four methods (other than *smartid*), gave fairly similar scores for the two monocytes subtypes (FCGR3A and CD14 monocytes), as well as for NK and CD8 T cells (**Figure 4A-F**). This suggests the difficulty for these four methods to discern these cell types, resulting in an overall lower AUPRC across all cell types (in **Figure 4G** and **Supp Figure 6**). In contrast, *smartid* consistently gave the highest AUPRC, particularly with the use of labels, highlighting its superior ability to discern similar cell types (Error! Reference source not found.**G** and **Supp Figure 6**).

Notably, even without labels, *smartid* still performed better than the other methods. This was particularly evident for FCGR3A monocytes and NK cells (**Supp Figure 6**), the smallest populations in the dataset (**Supp Figure 5H)**. These results demonstrated the power of *smartid* to accurately discriminate different cell types, particularly in the context of small or rare cell populations within imbalanced data.

### *smartid*-based score improves cell population separation

As *smartid* assigns a score to each gene in each cell (using equation (3.2), outlined in **Methods**), it can be used as a data transformation method for clustering. Compared to the traditional data transformation method (normalized log-counts), clustering based on the *smartid* score can improve the separation of cell populations. The dimensionality reduction analyses were conducted using the log-counts from *scater* (22) and the *smartid* scores, and the results were compared using t-distributed Stochastic Neighbour Embedding (tSNE) plots. The clustering based on the *smartid* score is clearly able to separate the two monocyte subtypes (FCGR3A and CD14 monocytes), as well as NK and CD8 T cells (**Figure 4H**). On the other hand, the clustering based on log-counts shows overlapping clusters of these similar subtypes, suggesting poorer separation (**Figure 4I)**. Furthermore, CD14 monocytes were split into two distinct subclusters by *smartid* (**Figure 4H**), indicating the presence of potential subtypes within CD14 cells. This finding highlights the potential of *smartid* to discover novel subtypes and expand the exploration of the functional classification in monocytes. In addition, it was noted that using only the TF-IAE score without the IDF factor was able to achieve comparable performance to the full TF-IDF-IAE score (**Figure 4J**), suggesting the ability of TF-IAE score to separate closely similar cell types.

### *smartid*-identified markers uncover cell-type-associated biological relevance on single cell data

To assess the application of *smartid* to elucidate mechanisms of biological significance, we applied it to a public scRNA-seq COVID19 patient dataset (‘scCOVID19’) (23). The marker genes for each cell type were identified from the top 100 genes ranked by *smartid* importance. These genes are able to identify the corresponding cell type and clearly separate it from others (**Figure 4K**). This was maintained when using *scran*-normalized logcounts (**Supp Figure 7**). To assess if these marker genes accurately reflect the biological processes associated with their respective cell types, we performed geneset enrichment analysis (GSEA) and visualization on the main biological themes using the R/Bioconductor package *vissE* (24).

The result for T-NK-ILC cell marker genes (a cell type in the data label containing all T/NK/ILC-like cells) was shown, where the network of the enriched gene-sets showed the presence of several large distinct clusters (**Figure 4L**), each representing a main biological theme. The word cloud for each cluster identifies the major biological themes represented in the gene-sets from that cluster (**Figure 4M)**. Gene-set clusters 2 and 3 show a significant enrichment of gene-sets associated with the apoptosis and caspase pathways. In contrast, clusters 1, 4 and 6 present a high activity of the gene-sets related to immune cells, particularly T cells and leukocytes. These findings are consistent with the biological roles of T-NK-ILC cells and support the involvement of these cells in the broader inflammatory responses in COVID19 patients (25). Results for other cell types are shown in **Supp Figure 8**. These results demonstrate that the *smartid*-derived marker genes are not only effective in distinguishing between cell populations, but also provide valuable insights into the biological processes, pathways and mechanisms for these populations.

### *smartid* inherently handles the batch effect and accurately identifies markers

*smartid* is able to identify cell-line-specific marker genes across platform batches without prior batch correction of the data. To assess how well *smartid* is able to account for batch effects in the data, we applied *smartid* on a well-designed experiment dataset (‘scdata’) generated by Tian, L. et al. (26). This data includes 1401 cells from 3 different cell lines (H1975, H2228 and HCC827), sequenced at three different platforms (10X Chromium, CEL-Seq2 and Drop-Seq) (**Figure 5A**). A standard data preprocessing workflow using *scater* (22) was applied, with all 3 datasets from the different platforms subsequently merged.

**Figure 5.**
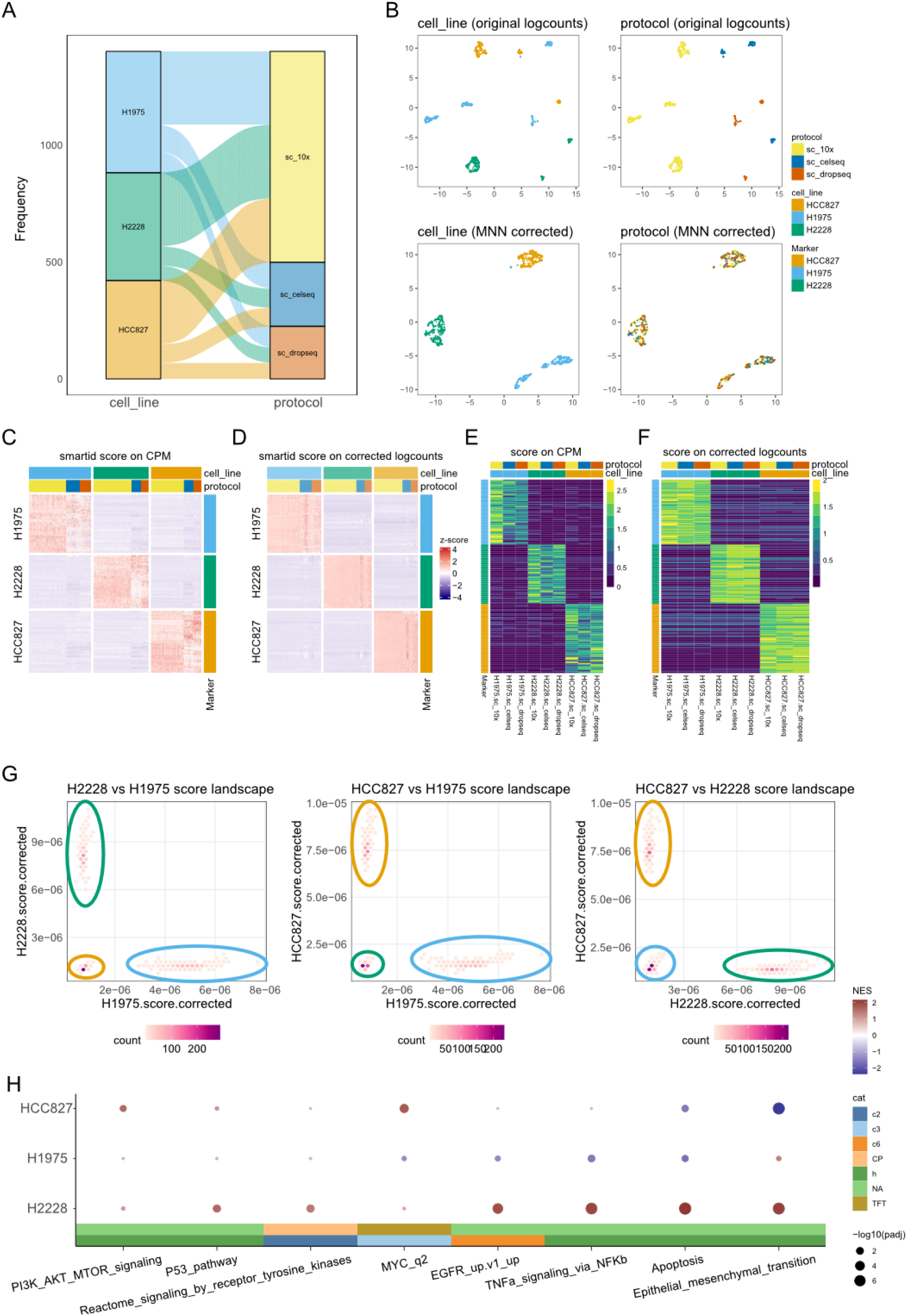
Cluster marker expression and score on scdata. (A) Sankey diagram depicting the cell line composition and platform protocol distribution. (B) UMAP of ‘scdata’ (26) coloured by cell line (left) and platform protocol (right), UMAP was performed on original logcounts (uncorrected, top) and MNN corrected logcounts (bottom). (C)-(D) Heatmap of scaled *smartid* score on CPM (C) and MNN corrected logcounts (D) at single cell level, using the top 50 *smartid*-derived marker genes for each cell line. (E)-(F) Heatmap of scaled average *smartid* score on CPM (E) and MNN corrected logcounts (F) at group level, using the top 50 *smartid*-derived marker genes for each cell line. (G) Density landscape of *smartid* score of cell lines, circled by cell line. (H) GSEA dotplot of interesting gene sets based on genes ranked by *smartid* importance.

Firstly, we assessed the UMAPs of the logcounts data processed by *scater* where a strong batch effect can be seen across different platforms. Applying the commonly used batch correction MNN (**Figure 5B)**, the batch can be seen to be corrected where cells now cluster primarily by cell line. In the following assessments, we use this MNN corrected data as a proxy for batch corrected data to assess the performance of *smartid*’s internal batch correction capabilities.

To evaluate *smartid*’s capability to handle strong batch effects without prior batch correction, the unnormalized CPM was used in conjunction with platform protocol as the batch factor to be regressed out. In summary, *smartid* identified 246 marker genes for H1975, 407 marker genes for H2228, and 210 for HCC827. Looking at the *smartid* score of normalised CPM expression patterns (without batch adjustments) of the top 50 *smartid* identified marker genes for each cell line (**Figure 5C**), significant platform-dependent variations in marker gene expressions can be observed, particularly notable differences can be seen between 10X Chromium and the other platforms. However, platform-specific expression patterns (**Supp Figure 9**) suggest that the identified marker genes maintained distinct expression profile within each platform, implying that while the uncorrected expression data exhibited platform-specific bias, the marker selection itself remained unbiased. This was further validated by examining the gene expression pattern scores of the marker genes using MNN corrected logcounts (**Figure 5D**), where minimal batch variation was observed, once again implying that the *smartid* identified markers are batch independent. Notably, the batch effect correction achieved by *smartid* scoring (via marker expressions on MNN corrected logcounts in **Figure 5D**) appears to be better than when using just MNN-correction (**Supp Figure 10**). The result is more evident at the individual marker gene expression level stratifying cells by cell line and platform (**Figure 5E & F**), where less platform batch effect was observed in batch corrected data.

We then computed the aggregate gene set score of each cell line’s markers for each cell. All three cell line’s marker genes demonstrated excellent discriminatory capacity, with each marker list easily able to isolate the corresponding cell line (**Supp Figure 11A**). These marker lists have exceptionally high specificity and sensitivity scores (area under the precision-recall curve, AUPRC) (**Supp Figure 11B**). Projecting the cells onto *smartid* score landscape, any pair cell line specific scores was sufficient to distinguish the three different cell lines independent of batch (**Figure 5G**).

Lastly, we investigated the functional significance of the markers genes identified for each cell line using pathway analysis (GSEA, **Figure 5H)** with results consistent with the known pathways for each cell line. Namely, H2228 has high activity of EGFR and TP53 pathways (27,28), H1975 has significant enrichment of EMT related genes (28) and HCC827 shows significantly higher activity of AKT and MYC related pathways (29).

These results and validations highlighted the robust nature of the *smartid* approach in correcting batch effects while preserving the biological signals in the data.

### *smartid* identified markers recapitulates the spatial cellular structure in the lymph node

To evaluate *smartid*’s utility in analysing image-based spatial transcriptomics data, it was applied to a single-cell spatial transcriptomics lymph node dataset generated using the Xenium platform (see **methods**), comprising 156,449 cells with 7,932 detectable genes. Visualizing the provided labels (from 10X Genomics) on the spatial coordinates, the follicular architecture of B cells, surrounded by endothelial cells can be clearly seen (**Figure 6A, Supp Figure 12A&C**). To assess *smartid*’s performance under rare cell type conditions, we randomly down-sampled both B cells and endothelial cells to each represent 0.1% of the total cell population, applied *smartid* on the raw counts and identify markers for the cell types.

**Figure 6.**
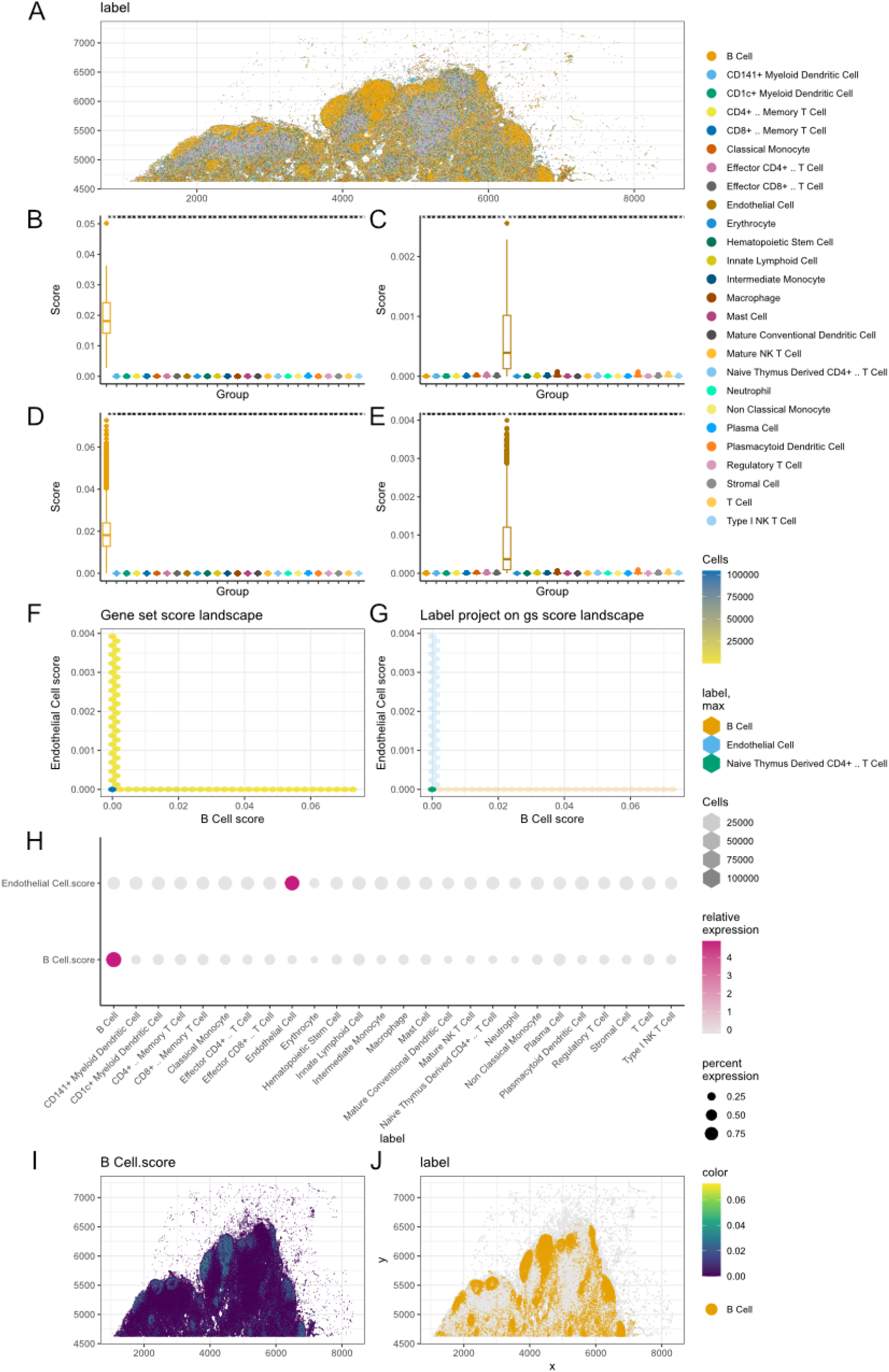
Marker score on spatial data. (A) Spatial map of cells stratified using the provided cell type label. (B-E) *smartid*-scores of B cell (B, D) and endothelial cell (C, E) markers across the cell type groups. Scores are either calculated on the down-sampled sub-dataset (B&C) or on the original dataset (D&E). (F-G) Endothelial cell score (y axis) plotted against B cells score (x axis) with hex bins of cells stratified based on density of cell count (F) and cell type (G). (H) Dotplot of marker score across cell types. (I) B cell marker score mapped on spatial slide. (J) B cell label mapped on spatial slide.

Scoring the *smartid*-identified markers across the label groups provided in the down-sampled sub-dataset, the respective scores can clearly distinguish either B cells or endothelial cells from all other cell types (**Figure 6B & C**). These markers, when scored across the groups in the original dataset remained highly discriminative for either cell type (**Figure 6D-H**), demonstrating the robustness of *smartid* markers. By visualising the *smartid*-derived B cell marker scores on the spatial coordinates, the characteristic lymphoid structures with B cell-dominated follicles (**Figure 6I**) can be seen, which strongly correlates with provided cell type labels (**Figure 6J, Supp Figure 12D**). Similarly, the endothelial cell marker scores also correlate well with the endothelial cell labels and noted to be localised to the boundary surrounding the B cells (**Supp Figure 12C&E**), recapitulating the structural organization of lymph node tissue.

We then visualized some of the identified B cell marker genes on the spatial coordinates, with spatial expression pattern highly correlating with that of the B cell labels (**Supp Figure 13, Figure 6A**). Notably, all five identified markers are well-known B cell surface proteins, including MS4A1 (CD20), CD79A and CD19 which are canonical markers of follicular B cells (30), CD22 and TNFRSF13C (BAFF-R) are characteristically expressed by marginal zone B cells(31). This underscores *smartid*’s capacity to identify functionally relevant markers reflective of the biology of interest. Till this end, *smartid* can be utilised to analyse single-cell spatial data and in this particular dataset, able to accurately identify highly specific marker genes for B cells and endothelial cells from raw counts and under highly imbalanced conditions.

## Discussion

Marker gene identification is a crucial step in single-cell data analysis and remains a major research focus. However, many existing methods, such as *Seurat* (14) or *scran* (15), are unable to handle the challenge of imbalanced data, which can bias marker gene identification. To address this issue, we developed *smartid*, an R/Bioconductor software package designed to identify marker genes from imbalanced data, with a particular focus on rare cell populations.

Having benchmarked *smartid* against 4 popular methods (*Seurat* (14), *Monocle3* (7), *scran* (15) and *CelliD* (16)) in this work, *smartid* not only accurately identifies effective marker genes from imbalanced datasets but also outperforms existing methods in this task. This stems primarily from the ability of *smartid* to avoid misleading marker genes. This improvement in accuracy aims to reduce or prevent the unnecessary waste of time and resources on pursuing experimental validation of potentially inaccurate results and to also improve the overall reliability and credibility of subsequent analyses and applications. Performance on biological datasets further demonstrates the advantages of *smartid*, especially in distinguishing between similar or rare cell types.

As a gene set scoring method, *smartid* shows superior discriminatory ability compared to existing methods, effectively characterizing cell type heterogeneity and improving clustering in low-dimensional spaces. Notably, even when using the TF-IAE score alone, without the IDF component, cell population separation remains clear and improved. This flexibility allows researchers to tailor the scoring method to their specific dataset and research purposes, potentially saving computational time and resources. The framework of *smartid* supports such customizations, offering users a range of options to customize the scoring method by combining different combinations of TF-IDF-IAE in their analyses. In addition, *smartid* successfully identifies the marker genes that are related to the biological processes of the corresponding cell types, which is valuable in elucidating cellular functions and molecular mechanisms.

Our comprehensive analysis also demonstrates that *smartid* effectively addresses one of the more challenging issues of scRNA-seq analysis, i.e. the integration and interpretation of data across different batches (32). The in-built capability of *smartid* to correct for known batch and identify population-specific markers, even in the presence of substantial batch effects, represents a significant advancement in single-cell data analysis methodology. These findings have several important implications for the field. First, it suggests that *smartid* could serve as a valuable tool for cross-platform data integration, potentially enabling more robust meta-analyses of single-cell datasets. Second, the method’s ability to identify markers effectively without prior batch correction could simplify analytical workflows and reduce the risk of introducing artifacts through preprocessing steps. Finally, the strong discriminatory power of *smartid*-derived markers indicates potential applications in developing robust cell-type classification systems that are resilient to technical variations.

With the recent emergent of spatial biology and profiling of molecular and cellular structures in tissues, it’s one of the important utilities that *smartid* can have substantial impact on. In fact, our analysis on single cell spatial lymph node suggests some interesting insights about *smartid’s* application to this newly emerging field. Importantly, *smartid* was able to successfully identify specific B and endothelial cell markers when the B cells and endothelial cells composition was adjusted to just 0.1% of all cells, simulating an imbalance dataset condition. In the context of spatial data analysis, it is already a challenging endeavour to work with such small number of cells (<100 cells after down-sampling for B or endothelial cells) in spatial data. However, it is relevant because many important cell types are naturally low in numbers in tissues, particular so in spatial dataset which is inherently sparse and especially so in tumour tissues where immune cells are rare. Spatial data provided the crucial validation not usually afforded in conventional single cell studies where only clusters of cells with marker genes providing the key validation avenue. Instead, spatial data provided the direct validation based on locality and spatial organisation of cell types. In this study, the observation of the well-established B-cell rich follicles/germinal centres as well as the surrounding endothelial tissues provided a visual and direct validation of the biology, which is very powerful way to ascertain the effectiveness of *smartid* as a marker identification tool. Examination of individual marker gene further confirmed the validity of markers in practical applications, such as the utility in flow cytometry.

In our spatial analysis investigations, what really caught our attention was the fact that the markers from the down-sampled data still works well on the full dataset. This suggests *smartid* is discovering fundamental characteristics of these cell types that hold up across different conditions, which could be useful for translating findings between studies or even into clinical settings.

There are a few limitations associated with *smartid*. Firstly, it currently uses a fixed threshold based on the total number of genes to identify the marker genes. This same threshold is applied to all different cell types within the same dataset. In reality, different cell types will have different importance and thus different distributions and weightage, pointing towards having cell type-specific thresholds to further improve feature selection. This will require further development and optimization which has been factored into the future works. Secondly, when using *smartid* as a scoring method, it is currently only suitable for comparing scores of different cells within a specific gene panel, and not application to compare scores of the same cell with different set of gene. An improved normalization will be needed for *smartid* to be able to address this limitation. Thirdly, with the label-based scores significantly performing better the unlabelled scores, improvements to the scoring algorithm for label-free data will be need in future development to reduce the package’s dependence on data labels.

Finally, future studies could explore the application of *smartid* to more complex experimental designs, including time-series data, spatial data and multi-omics data. Investigating the method’s performance across different omics technologies and experimental protocols could further validate its broad applicability.

## Methods

### Modifying TF-IDF for scRNAseq

By considering cells as ‘documents’ and genes as ‘terms’, TF-IDF is adapted for scRNA-seq data (as in equations (1–5)). Employing the concept of TF-IDF, each gene is assigned a weight for each cell, representing the ‘importance’ of that gene to the cell. The TF gives more weight to the genes with a higher expression in the cells and the IDF gives more weight to the genes with higher specificity, particularly those from a small or rare population. However, as scRNA-seq data is a lot sparser than text documents, this may lead to a large IDF value when there are no cells expressing a particular gene. In another situation, for genes expressed in all cells with significantly different expression levels in many cell types, the IDF will tend to be 1, making it hard to separate the specific marker genes from the stably expressed genes. To visualize these cases, we generated a simple data example with 5 main categories of genes from 3 cell types (**Supp Figure 1**).

The genes in red and yellow are expressed across all cells. However, the red gene is significantly more expressed in Type 1 cells, whereas the yellow gene is specifically expressed in Type 3 cells. Both the light blue and dark blue genes show stable expression across all cells: the light blue gene is highly expressed and stable, while the dark blue gene is consistently low or absent. The green gene is uniquely expressed in Type 2 cells and shows minimal expression in other cell types.

Due to their global expression, both the yellow and light blue genes will have an IDF of 1, resulting in similar TF-IDF values in Type 3 cells. This similarity makes it difficult for traditional TF-IDF score to distinguish between these two genes. In contrast, the dark blue gene, which is expressed at very low levels in most cells, has a high IDF value. This can lead to a disproportionately high TF-IDF score in cells where the dark blue gene is only marginally expressed, potentially distorting the gene selection process.

Consequently, the traditional TF-IDF score (equation (1)) is unable to identify three of the five categories in its current form. To improve the discriminative power of the score in equation (2), we derived a new factor IAE (Inverse Average Expression) modified from the IDF (equation (5). By replacing *M*_*i*_ with *G*_*i*_, the IAE can assign a higher weight to those genes with high expression in small populations having similar TF and IDF values. Specifically, when comparing the TF-IDF-IAE scores for the yellow and blue genes in Type 3 cells in **Supp Figure 1**, the IAE values are greater for the yellow gene relative to the light blue gene within Type 3 cells. This enables these genes to be distinguished, a task that the traditional TF-IDF score fails to achieve. Using this approach, *smartid* works on both unfiltered raw counts and processed data (e.g. CPM and RPKM).

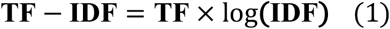

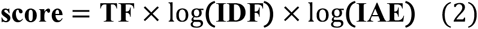

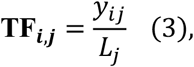

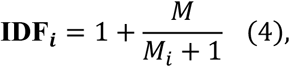

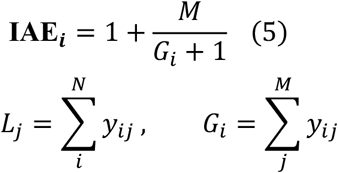

where *y*_*ij*_ is the count of gene *i* in cell *j, L*_*j*_ is the library size of cell *j, G*_*i*_ is the total counts of gene *i* across all cells, *N* is the number of genes, *M* is the total number of cells and *M*_*i*_ is the number of cells containing gene *i* with count > 0 (#{*j*: *y*_*ij*_ > 0}).

#### Scoring Unlabelled Data

*smartid* can be used to score unlabelled data using the standard IDF and IAE (in equations (4–5)). However, in cases where certain genes are lowly expressed across all cells (e.g. the dark blue gene in **Supp Figure 1**), the standard IAE may be disproportionately large, resulting in an incorrectly similar or even higher score than the highly expressed marker genes within the cell type. To resolve this, we included the standard deviation (SD) term into the IDF and the IAE in equations (6–7). Higher weight is added to those highly variable genes across all cells. Since the TF ranges from 0 to 1, the corresponding range for its variance is also between 0 and 1.

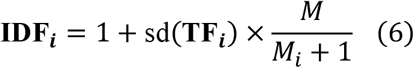

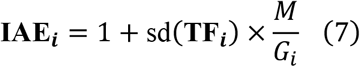

### Marker Identification by Score

#### Scoring Labelled Data

The SD version of the IDF and IAE can be used for unlabelled data. However, as marker gene selection is always performed on labelled data, we modified the IDF and IAE to incorporate label information as shown in equations (8–9). By introducing *K* classes, the IDF_*ijk*_ for gene *i* is no longer the same for all cells, but with a specific value for cell *j* in class *k*. This facilitates distinguishing genes with the same *M*_*i*_ but different *M*_*i,k*_, thereby enabling the assignment of different weights to gene *i* depending on class *k*. This new score_*ijk*_, using the labelled IDF_*ijk*_ and IAE_*ijk*_, allows for a more accurate estimation of the importance of the gene for the rare cell populations during feature selection, reducing the bias towards the dominant cell populations.

For gene *i* and cell *j* in class *k*:

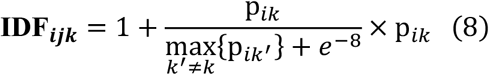

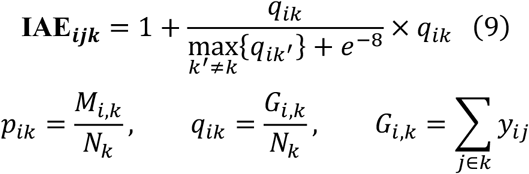

where *M*_*i,k*_ is the number of cells in class *k* containing gene *i, G*_*i,k*_ is the total counts of gene *i* across cells in class *k, N*_*k*_ is the number of cells in class *k*.

#### Scaling and Transformation

Each gene has a different range in terms of expression, which biases the feature selection towards those with large numerical ranges. Therefore, in labelled data, scaling is needed for the labelled score_*ijk*_ before the feature selection. However, applying the standard scaling methods on imbalanced data, such as min-max normalization or z-score normalization, may inadvertently modify the relative importance of minority/rare and majority/dominant classes, resulting in reduced distinguishability between them. To address this, we performed a weighted mean, where cells from rarer populations are assigned higher weights, ensuring that each population/class contributes equally at the class level (in equations (10–11)). With the weighted mean and the pooled SD *s*_*pooled*_, more robust estimates of the scaled score *z*_*ijk*_ can be obtained for labelled data, mitigating the effect of class imbalance.

For gene *i* and cell *j* in class *k*:

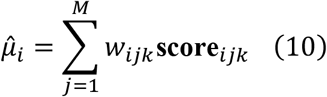

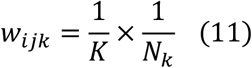

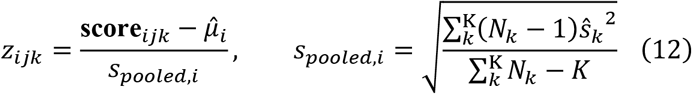

where *w*_*ijk*_ is the weight of gene *i* for cell *j* in class *k, K* is the total number of classes, 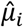 is the weighted mean of the score for gene *i, z*_ijk_ is the scaled score, *s*_*pooled,i*_ is the pooled SD of gene *i* and ŝ_*k*_ is the SD of gene *i* within class k.

The mean of the scaled score *z*_*ijk*_ for gene *i* across all cells in class *k* was estimated as 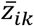, then the importance (*I*_*ik*_) of it was obtained by calculating the differences between the 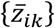 as described in equation (13). For each class, the original importances {*I*_*ik*_} of all genes were transformed using a softmax transformation function *g*(∙) to produce the transformed importances {*SI*_*ik*_ }, as shown in equation (14). This normalized exponential transformation constrained the importance within the range [0,1] and summed to 1.

For gene *i* across all cells in class *k*:

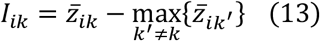

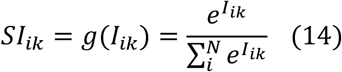

where *I*_*ik*_ is the original importance of gene *i* across all cells in class *k, SI*_*ik*_ is the transformed importance of *I*_*ik*_ and *g*(∙) is the softmax transformation function.

#### Batch Effect Correction

To mitigate the potential impact of batch effects on the data, we implemented a statistical framework based on the assumption that the scaled expression scores follow a Gaussian distribution. Specifically, we used a GLM to estimate the mean of the scaled score *z*_*ijk*_, incorporating the batch factor as a fixed effect in the model formulation to partition the variance attributable to class and batch (equations (15–16)). This approach allows for robust marker detection while accounting for systematic batch-to-batch variation in the underlying data structure.

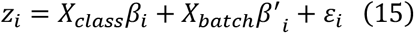

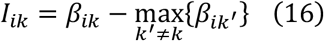

where *z*_*i*_ is the scaled score of gene *i* across all cells, *X*_*class*_ and *X*_*batch*_ are design matrices for the cell classes and batches, *β*_*i*_ and *β*′_*i*_ represent the fixed effects for the corresponding class and batch, 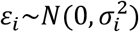 represents the remaining random error.

#### Feature Selection

we then modelled the transformed importance *SI*_*ik*_ in class *k* as a Gaussian finite mixture, which can accommodate different numbers of components. we used the R/CRAN package *mclust* (33) here to estimate the probability density function for the potential components. For *H* components of class *k*, the mixture model takes the form:

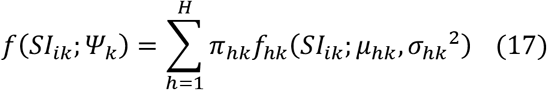

where *Ψ*_*k*_ are the parameters of the mixture model (*π*_*hk*_, *μ*_*hk*_, *σ*_*hk*_^2^), *μ*_*hk*_ is the mean and *σ*_*hk*_^2^ is the variance of the Gaussian model for component *h* of class *k, f*_*hk*_ is the density function of component *h, π*_*hk*_ is the mixing probability 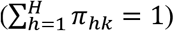.

For each fixed *H*, the unknown parameters *Ψ*_*k*_ were estimated using the EM algorithm (12). Scanning across a range of potential *H* (2 to 9), the best number of mixing components for GMM was selected by maximizing the Bayesian Information Criterion (BIC), where BIC penalizes more heavily with increasing number of components. The component with highest *μ*_*hk*_ will be considered as the density function of the marker genes, and the genes with probability over the threshold (0.99 by default) will be selected as final marker genes.

All statistical analyses were performed using R (version ≥ 4.4.0) and Bioconductor (version ≥ 3.19). We have produced an R/Bioconductor package, *smartid*, to implement this method, and have included the visualization functions to plot marker genes and score performance (using *ggplot2* (34)).

### Benchmarking

The performance in marker gene identification of some current methods was compared to *smartid*, including *Seurat* (14), *Monocle3* (7), *scran* (15) and *CelliD* (16). The accuracy was evaluated by computing the F1 score (equation (20)), recall ratio (equation (19)) and false discoveries based on a negative control group, while the efficiency was assessed by calculating the computational running time from the start to finish.

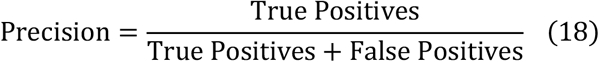

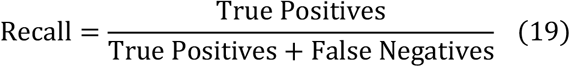

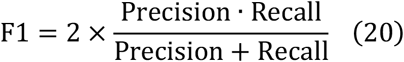

For gene-set scoring, *smartid* was compared with *UCell* (18), *GSVA* (19), *ssGSEA* (20) and *AUCell* (21). The area under the precision-recall curve (AUPRC) (using *precrec* (35)) of the score using different methods was calculated by comparing the score against the binarized cell label for each cell type (e.g., grouped as either ‘NK’ and ‘non-NK’ for the NK cell marker). All scores were computed using the marker genes identified by *smartid* from the dataset ‘pbmc3k.final’ (17). Default parameters were used for all the scoring methods.

### Gene-set enrichment analysis (GSEA)

GSEA was conducted using *clusterProfiler* (36) on the gene-sets under categories: ‘H’ (hallmark gene sets), ‘C2’ (curated gene sets), ‘C5’ (ontology gene sets) and ‘C7’ (immunologic signature gene sets), from MSigDB (37). The marker genes used in GSEA were identified from the COVID19 cell atlas (24) for each cell type using *smartid*. Enrichment results were further analysed and visualized using *vissE* (24) to explore the biological processes involved for each cell type.

## Datasets

### Simulated datasets

Simulated scRNA-seq raw count data was generated using *splatter* (13), consisting of 1,000 or 10,000 genes with 3,000 or 10,000 cells. The cells were divided into four cell groups, named Groups 1 to 4. Each group comprised 10%, 20%, 30% and 40% of the total population, respectively, to examine the performance of the methods on groups with different abundances. For further benchmarking on imbalanced data, each group comprised 1%, 20%, 30% and 49% of the total cell population (10,000 cells). Groups 1 to 3 each had 2% of all genes being differentially expressed genes (DEGs), while Group 4 with no DEGs was used as a negative control group. The logFC of the DEG was sampled from a normal distribution with a mean of 0.5 and SD of 0.4.

### Real-life datasets

The raw count data of the ‘pbmc3k.final’ dataset was obtained from *SeuratData* (17) to compare the performance between the methods on real-life biological data. Dendritic and platelet cells were excluded from ‘pbmc3k.final’ for the analysis, as there are less than 50 cells in each cell type. we also used the raw count data from the COVID19 Cell Atlas (23) (denoted as ‘scCOVID19’) to identify the marker genes of B/plasma cells, epithelial cells, macrophages, myeloid cells and T-NK-ILC cells using *smartid*. All the data was processed with default parameters using *smartid*.

The scRNA-seq data of 3 non-small cell lung cancer (NSCLC) cell lines sequenced at 3 platforms (26) (denoted as ‘scdata’) was used to evaluate the ability of *smartid* to deal with batch effect. To better handle the strong batch effect, *smartid* was applied on the unnormalized CPM values rather than raw counts. For downstream validation of the derived marker performance, we utilized log-transformed count data (denoted as ‘logcounts’) that had undergone mutual nearest neighbour (MNN) batch correction as implemented in *batchelor* (38).

The Xenium spatial transcriptomics dataset from human lymph node tissue was obtained from 10X Genomic (https://www.10xgenomics.com/datasets/preview-data-xenium-prime-gene-expression). The dataset contains spatially resolved single-cell expression profiles for 7,932 genes measured using the 10x Genomics Xenium platform. Raw gene counts were used directly as input to *smartid* without normalization, scaling, or other preprocessing steps to evaluate the method’s performance on unprocessed data. To simulate rare cell type detection scenarios, B cells and endothelial cells were randomly down-sampled to represent 0.1% of the total cell population while maintaining the original proportions of all other cell types. *smartid* was applied to the down-sampled dataset to identify marker genes specific to B cells and endothelial cells. The identified markers were then used to compute cell type-specific scores for all cells in both the down-sampled and original dataset.

## Data availability

All datasets underlying this article are available from public data repository or publications as described in **Table 2**.

**Table 2.**
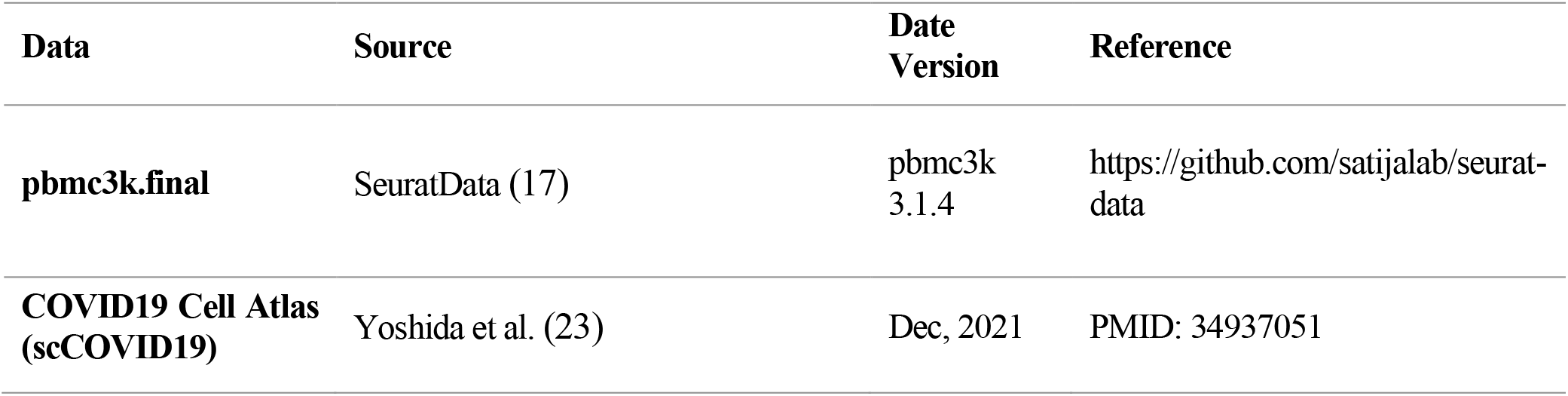
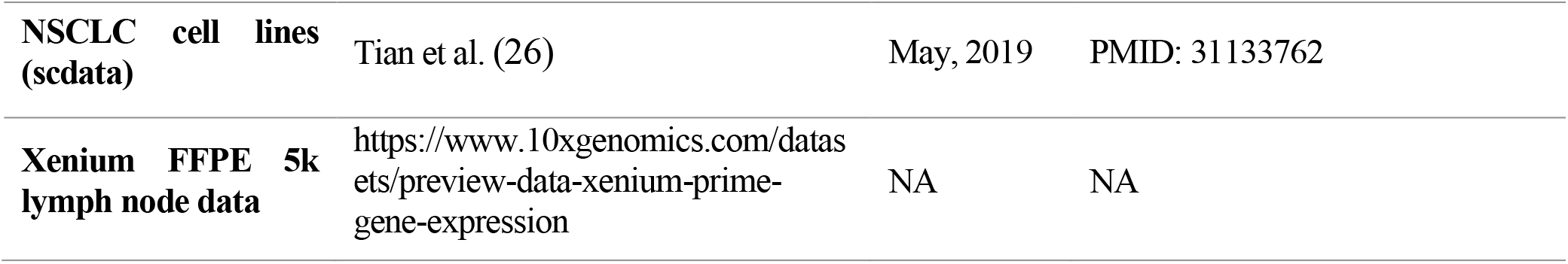
Datasets used in this study.

## Code availability

The code used for performing the described analyses is available from the corresponding author upon request. The *smartid* package is publicly available on Bioconductor (version ≥ 3.19) at https://bioconductor.org/packages/release/bioc/html/smartid.html. A comprehensive guide introduction can be found at https://davislaboratory.github.io/smartid/articles/smartid_Demo.html.

## Author contributions

J.C.: Conceptualization, Methodology, Software development, Formal analysis, Visualization, Project administration, Writing-Original Draft

M.L.: Methodology, Writing-Review & Editing

D.D.B.: Analysis advice, Writing-Review

M.J.D.: Supervision, Conceptualization

A.T.P.: Study design input, Supervision, Writing-Review & Editing

C.W.T.: Supervision, Conceptualization, Project administration, Resources, Writing-Review & Editing

## Acknowledgements

J.C. and M.L. are supported by Melbourne Research Scholarships (MRS) from the University of Melbourne. C.W.T. is supported by the Australian Academy of Sciences (AAS): Regional Collaborations Programme COVID-19 Digital Grants scheme and the National Health and Medical Research Council Medical Research (NHMRC) Future Fund MRF203110. D.D.B. is supported by NHMRC Investigator grant GNT2034141. A.T.P. was supported by a NHMRC Investigator Grant (2026643) and funding from the Lorenzo and Pamela Galli Medical Research Trust. A.T.P. and J.C. were supported by an NHMRC Synergy Grant (2036149). WEHI acknowledges the Victorian State Government Operational Infrastructure Support; and the Independent Research Institutes Infrastructure Support Scheme of the Australian Government National Health and Medical Research Council to WEHI.

We would like to thank Prof. Terence P. Speed, Dr Lizhong Chen, Dr Givanna Putri and Dr Samuel Lee for reviewing and providing constructive feedback for this manuscript.

## Supplementary data

Supplementary data are available online.

## References

1. Jaitin, D.A., Kenigsberg, E., Keren-Shaul, H., Elefant, N., Paul, F., Zaretsky, I., Mildner, A., Cohen, N., Jung, S., Tanay, A. et al. (2014) Massively Parallel Single-Cell RNA-Seq for Marker-Free Decomposition of Tissues into Cell Types. Science, 343, 776–779.

2. Olguin-Martinez, E., Ruiz-Medina, B.E. and Licona-Limon, P. (2021) Tissue-Specific Molecular Markers and Heterogeneity in Type 2 Innate Lymphoid Cells. Front Immunol, 12, 757967.

3. Zhang, X., Lan, Y., Xu, J., Quan, F., Zhao, E., Deng, C., Luo, T., Xu, L., Liao, G., Yan, M. et al. (2019) CellMarker: a manually curated resource of cell markers in human and mouse. Nucleic Acids Res, 47, D721–D728.

4. Wang, N., Hoffman, E.P., Chen, L., Chen, L., Zhang, Z., Liu, C., Yu, G., Herrington, D.M., Clarke, R. and Wang, Y. (2016) Mathematical modelling of transcriptional heterogeneity identifies novel markers and subpopulations in complex tissues. Sci Rep, 6, 18909.

5. Liu, L.P., Wu, Q., Zhong, W.W., Chen, Y.P., Zhang, W.Y., Ren, H.L., Sun, L. and Sun, J.H. (2020) Microarray Analysis of Differential Gene Expression in Alzheimer’s Disease Identifies Potential Biomarkers with Diagnostic Value. Med Sci Monitor, 26.

6. Twomey, J.D., Brahme, N.N. and Zhang, B. (2017) Drug-biomarker co-development in oncology - 20 years and counting. Drug Resist Updat, 30, 48–62.

7. Trapnell, C., Cacchiarelli, D., Grimsby, J., Pokharel, P., Li, S.Q., Morse, M., Lennon, N.J., Livak, K.J., Mikkelsen, T.S. and Rinn, J.L. (2014) The dynamics and regulators of cell fate decisions are revealed by pseudotemporal ordering of single cells. Nature Biotechnology, 32, 381–U251.

8. Kharchenko, P.V., Silberstein, L. and Scadden, D.T. (2014) Bayesian approach to single-cell differential expression analysis. Nat Methods, 11, 740–742.

9. Mishra, A. and Vishwakarma, S. (2015) Analysis of TF-IDF Model and its Variant for Document Retrieval. Int Conf Comput Inte, 772–776.

10. Arnesia, P.D. and Madenda, S. (2012) Matching Images With Textual Document Using TFIDF Method. 2012 5th International Congress on Image and Signal Processing (Cisp), 1283–1289.

11. Aninditya, A., Hasibuan, M.A. and Sutoyo, E. (2019) Text Mining Approach Using TF-IDF and Naive Bayes for Classification of Exam Questions Based on Cognitive Level of Bloom’s Taxonomy. 2019 Ieee International Conference on Internet of Things and Intelligence System (Iotais), 112–117.

12. Moon, T.K. (1996) The expectation-maximization algorithm. Ieee Signal Proc Mag, 13, 47–60.

13. Zappia, L., Phipson, B. and Oshlack, A. (2017) Splatter: simulation of single-cell RNA sequencing data. Genome Biol, 18, 174.

14. Hao, Y., Stuart, T., Kowalski, M.H., Choudhary, S., Hoffman, P., Hartman, A., Srivastava, A., Molla, G., Madad, S., Fernandez-Granda, C. et al. (2024) Dictionary learning for integrative, multimodal and scalable single-cell analysis. Nat Biotechnol, 42, 293–304.

15. Lun, A.T., McCarthy, D.J. and Marioni, J.C. (2016) A step-by-step workflow for low-level analysis of single-cell RNA-seq data with Bioconductor. F1000Res, 5, 2122.

16. Cortal, A., Martignetti, L., Six, E. and Rausell, A. (2021) Gene signature extraction and cell identity recognition at the single-cell level with Cell-ID. Nat Biotechnol, 39, 1095–1102.

17. Lab, S. (2020).

18. Andreatta, M. and Carmona, S.J. (2021) UCell: Robust and scalable single-cell gene signature scoring. Comput Struct Biotec, 19, 3796–3798.

19. Hänzelmann, S., Castelo, R. and Guinney, J. (2013) GSVA: gene set variation analysis for microarray and RNA-Seq data. Bmc Bioinformatics, 14.

20. Barbie, D.A., Tamayo, P., Boehm, J.S., Kim, S.Y., Moody, S.E., Dunn, I.F., Schinzel, A.C., Sandy, P., Meylan, E., Scholl, C. et al. (2009) Systematic RNA interference reveals that oncogenic-driven cancers require TBK1. Nature, 462, 108–U122.

21. Aibar, S., Gonzalez-Blas, C.B., Moerman, T., Huynh-Thu, V.A., Imrichova, H., Hulselmans, G., Rambow, F., Marine, J.C., Geurts, P., Aerts, J. et al. (2017) SCENIC: single-cell regulatory network inference and clustering. Nat Methods, 14, 1083–1086.

22. McCarthy, D.J., Campbell, K.R., Lun, A.T.L. and Wills, Q.F. (2017) Scater: pre-processing, quality control, normalization and visualization of single-cell RNA-seq data in R. Bioinformatics, 33, 1179–1186.

23. Yoshida, M., Worlock, K.B., Huang, N., Lindeboom, R.G.H., Butler, C.R., Kumasaka, N., Dominguez Conde, C., Mamanova, L., Bolt, L., Richardson, L. et al. (2022) Local and systemic responses to SARS-CoV-2 infection in children and adults. Nature, 602, 321–327.

24. Bhuva, D.D., Tan, C.W., Liu, N., Whitfield, H.J., Papachristos, N., Lee, S.C., Kharbanda, M., Mohamed, A. and Davis, M.J. (2024) vissE: a versatile tool to identify and visualise higher-order molecular phenotypes from functional enrichment analysis. Bmc Bioinformatics, 25.

25. Silverstein, N.J., Wang, Y., Manickas-Hill, Z., Carbone, C., Dauphin, A., Boribong, B.P., Loiselle, M., Davis, J., Leonard, M.M., Kuri-Cervantes, L. et al. (2022) Innate lymphoid cells and COVID-19 severity in SARS-CoV-2 infection. Elife, 11.

26. Tian, L., Dong, X., Freytag, S., Le Cao, K.A., Su, S., JalalAbadi, A., Amann-Zalcenstein, D., Weber, T.S., Seidi, A., Jabbari, J.S. et al. (2019) Benchmarking single cell RNA-sequencing analysis pipelines using mixture control experiments. Nat Methods, 16, 479–487.

27. Voena, C., Di Giacomo, F., Panizza, E., D’Amico, L., Boccalatte, F.E., Pellegrino, E., Todaro, M., Recupero, D., Tabbo, F., Ambrogio, C. et al. (2013) The EGFR family members sustain the neoplastic phenotype of ALK+ lung adenocarcinoma via EGR1. Oncogenesis, 2, e43.

28. Gladilin, E., Ohse, S., Boerries, M., Busch, H., Xu, C., Schneider, M., Meister, M. and Eils, R. (2019) TGFbeta-induced cytoskeletal remodeling mediates elevation of cell stiffness and invasiveness in NSCLC. Sci Rep, 9, 7667.

29. Pilling, A.B., Kim, J., Estrada-Bernal, A., Zhou, Q., Le, A.T., Singleton, K.R., Heasley, L.E., Tan, A.C., DeGregori, J. and Doebele, R.C. (2018) ALK is a critical regulator of the MYC-signaling axis in ALK positive lung cancer. Oncotarget, 9, 8823.

30. Yao, L., Wang, J.T., Jayasinghe, R.G., O’Neal, J., Tsai, C.F., Rettig, M.P., Song, Y., Liu, R., Zhao, Y., Ibrahim, O.M. et al. (2023) Single-Cell Discovery and Multiomic Characterization of Therapeutic Targets in Multiple Myeloma. Cancer Res, 83, 1214–1233.

31. Muller-Winkler, J., Mitter, R., Rappe, J.C.F., Vanes, L., Schweighoffer, E., Mohammadi, H., Wack, A. and Tybulewicz, V.L.J. (2021) Critical requirement for BCR, BAFF, and BAFFR in memory B cell survival. J Exp Med, 218.

32. Hicks, S.C., Townes, F.W., Teng, M. and Irizarry, R.A. (2018) Missing data and technical variability in single-cell RNA-sequencing experiments. Biostatistics, 19, 562–578.

33. Scrucca, L., Fop, M., Murphy, T.B. and Raftery, A.E. (2016) mclust 5: Clustering, Classification and Density Estimation Using Gaussian Finite Mixture Models. R J, 8, 289–317.

34. Wickham, H. (2009) ggplot2: Elegant Graphics for Data Analysis. Use R, 1–212.

35. Saito, T. and Rehmsmeier, M. (2017) Precrec: fast and accurate precision-recall and ROC curve calculations in R. Bioinformatics, 33, 145–147.

36. Wu, T.Z., Hu, E.Q., Xu, S.B., Chen, M.J., Guo, P.F., Dai, Z.H., Feng, T.Z., Zhou, L., Tang, W.L., Zhan, L. et al. (2021) clusterProfiler 4.0: A universal enrichment tool for interpreting omics data. Innovation-Amsterdam, 2.

37. Liberzon, A., Birger, C., Thorvaldsdottir, H., Ghandi, M., Mesirov, J.P. and Tamayo, P. (2015) The Molecular Signatures Database Hallmark Gene Set Collection. Cell Syst, 1, 417–425.

38. Haghverdi, L., Lun, A.T.L., Morgan, M.D. and Marioni, J.C. (2018) Batch effects in single-cell RNA-sequencing data are corrected by matching mutual nearest neighbors. Nat Biotechnol, 36, 421–427.

