## Supplementary Figures for "Improved marker detection for rare population in single-cell transcriptomics through text mining-inspired scoring approach"

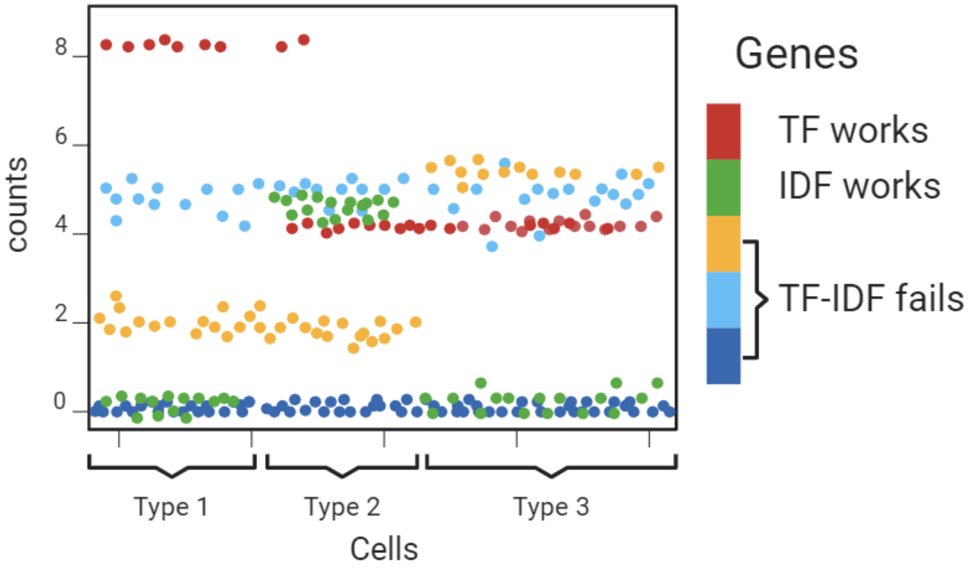


**Supp Figure 1. Gene expression profiles into five main categories.**

The red gene is highly expressed in Type 1 cells, the yellow gene is specific to Type 3 cells, and the green gene is unique to Type 2 cells. The light blue gene is stably high in all cells, while the dark blue gene is consistently low.

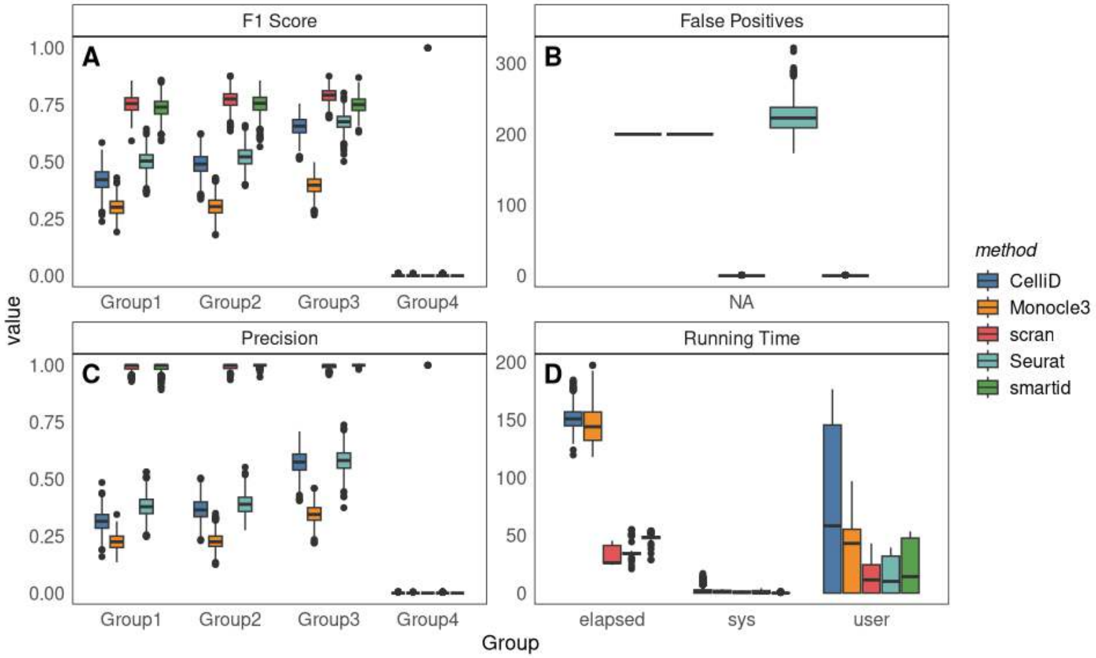


**Supp Figure 2. Performance of the evaluated methods on 1,000 simulated scRNA-seq datasets (3k cells).**

Benchmark results on simulation with 3k cells of 10k genes, each group consists of 10%, 20%, 30%, 40% cells (from Group 1 to 4). Boxplot of F1 score (A) and precision ratio (B) of identified marker genes across different groups using different methods. (C) Boxplot of false positives identified by different methods in negative control Group 4. (D) Boxplot of computational running time for marker gene identification using different methods, 'elapsed' (elapsed time) measures the total real time taken, ‘sys’ (system time) measures the amount of CPU time taken in system mode, ‘user’ (user time) measures the amount of CPU time taken in user mode. All boxes are coloured by methods and grouped by cell group labels. F1 score and precision was computed by comparing the identified marker genes with true DEGs in each group and false positives were computed by counting the identified false marker genes in Group 4 (which has no DEGs). Running time was captured for the whole marker gene identification process.

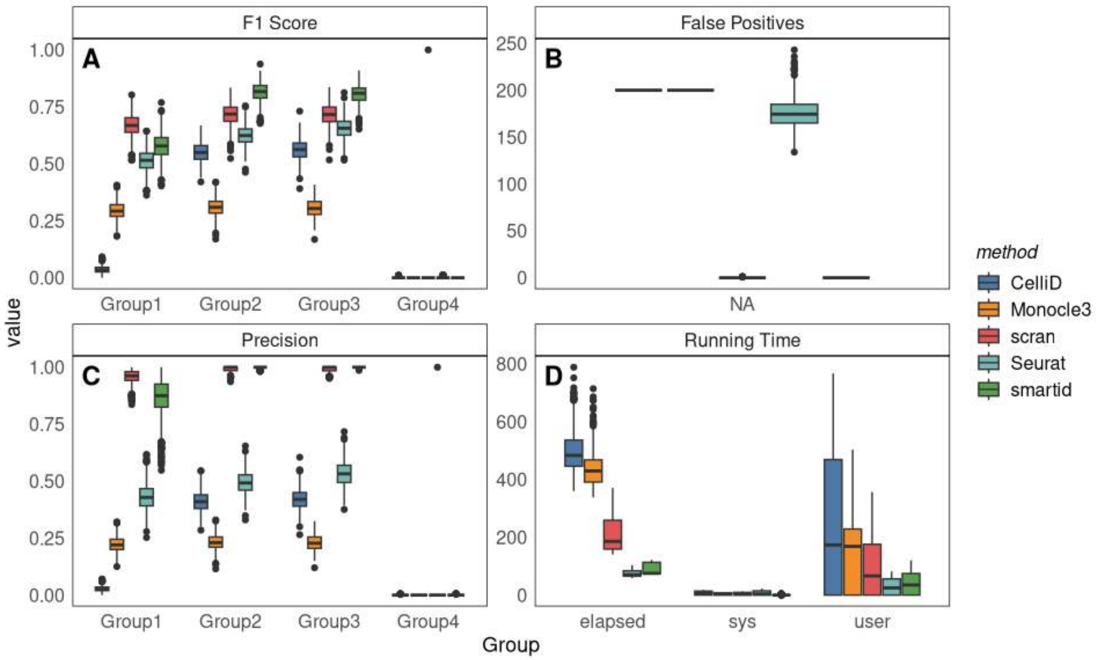


**Supp Figure 3. Performance of the evaluated methods on 1,000 simulated scRNA-seq datasets (10k cells).**

Benchmark results on simulation with 10k cells of 10k genes, each group consists of 1%, 20%, 30%, 49% cells (from Group 1 to 4). Boxplot of F1 score (A) and precision ratio (B) of identified marker genes across different groups using different methods. (C) Boxplot of false positives identified by different methods in negative control Group 4. (D) Boxplot of computational running time for marker gene identification using different methods, ‘elapsed’ (elapsed time) measures the total real time taken, ‘sys’ (system time) measures the amount of CPU time taken in system mode, ‘user’ (user time) measures the amount of CPU time taken in user mode. All boxes are coloured by methods and grouped by cell group labels. F1 score and precision was computed by comparing the identified marker genes with true DEGs in each group and false positives were computed by counting the identified false marker genes in Group 4 (which has no DEGs). Running time was captured for the whole marker gene identification process.

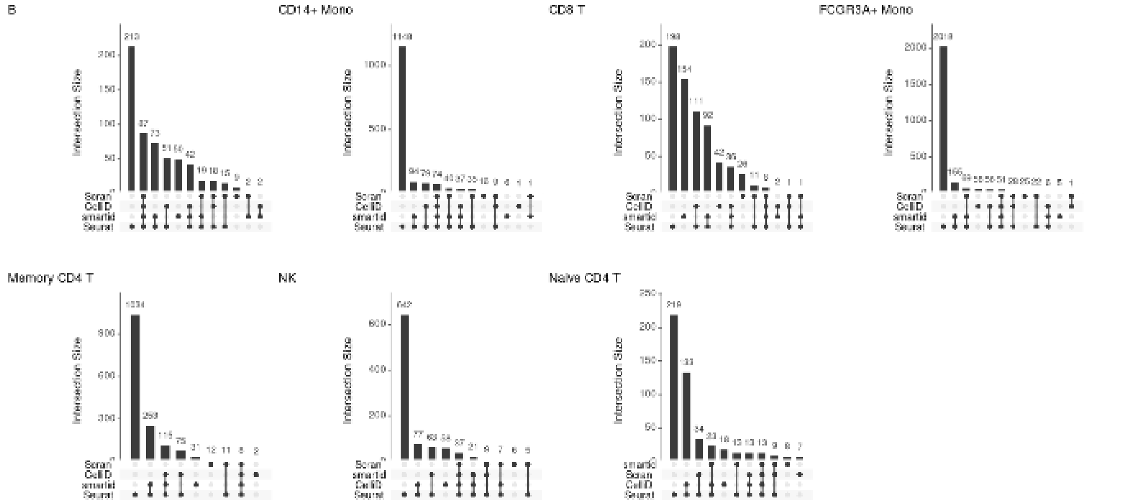


**Supp Figure 4. Overlap of marker genes identified by different methods for different cell types on pbmc3k.final dataset.**

The plot shows the intersections of marker genes identified by different methods for each cell type. Each bar represents the number of marker genes common to multiple methods.

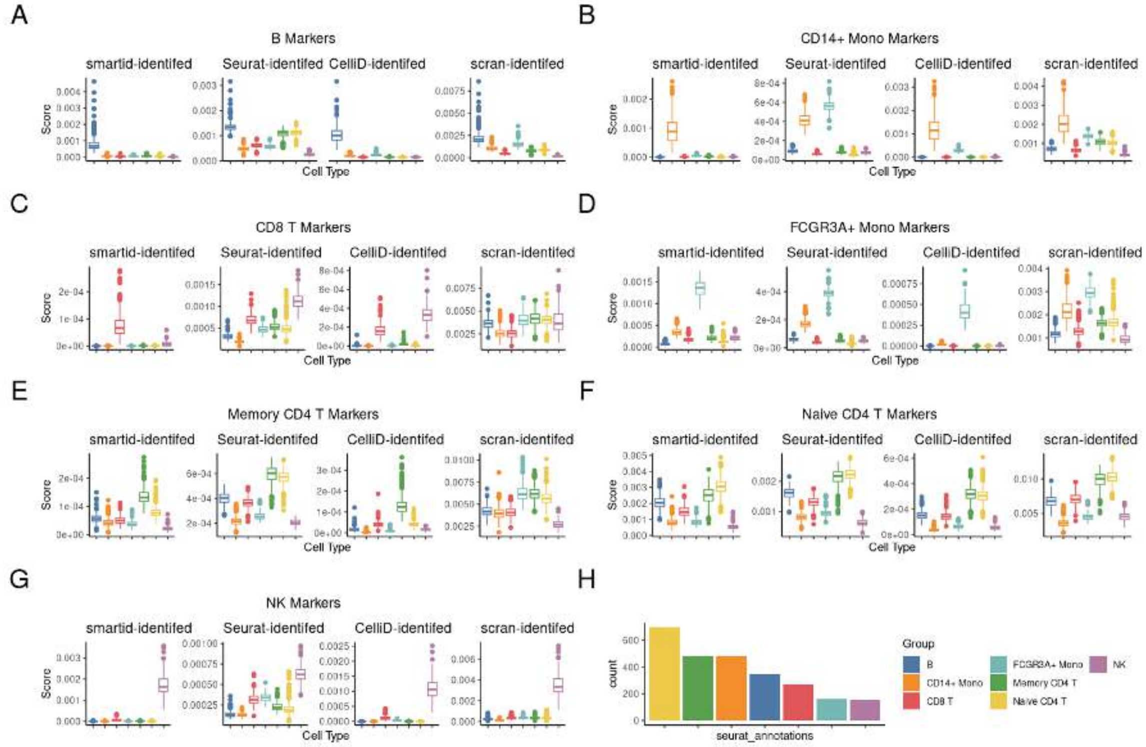


**Supp Figure 5. Boxplot of the score calculated by *smartid* using marker genes identified by different methods for each cell type on pbmc3k.fianl.**

(A)-(G) Boxplot of *smartid* score across different cell types by using the markers derived from different methods (from left to right: *smartid*, *Seurat*, *CelliD* and *scran*) for each cell type, where A-G are B cells, CD14+ monocytes, CD8+ T cells, FCGR3A+ monocytes, memory CD4+ T cells, naïve CD4+ T cells and NK cells in that order. (H) The bar plot of cell type proportions. All boxes and bars are coloured by cell type.

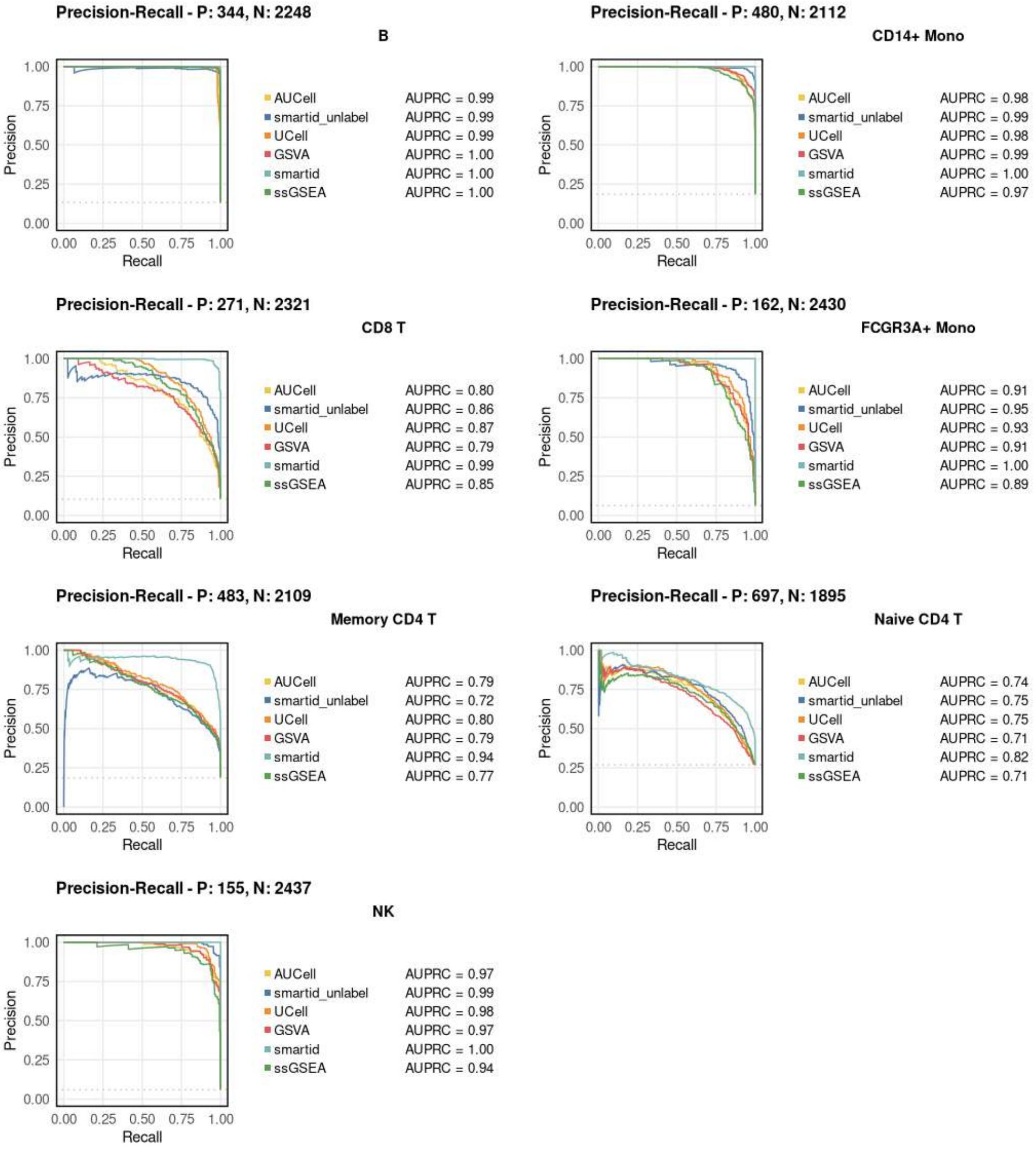


**Supp Figure 6. Precision-recall curves of different scoring methods using *smartid*-identified cell type marker genes.**

The x-axis represents the recall, the y-axis denotes the precision. AUPRC was computed by comparing the scores to the binarized labels for each cell type, curves are coloured by the different scoring methods.

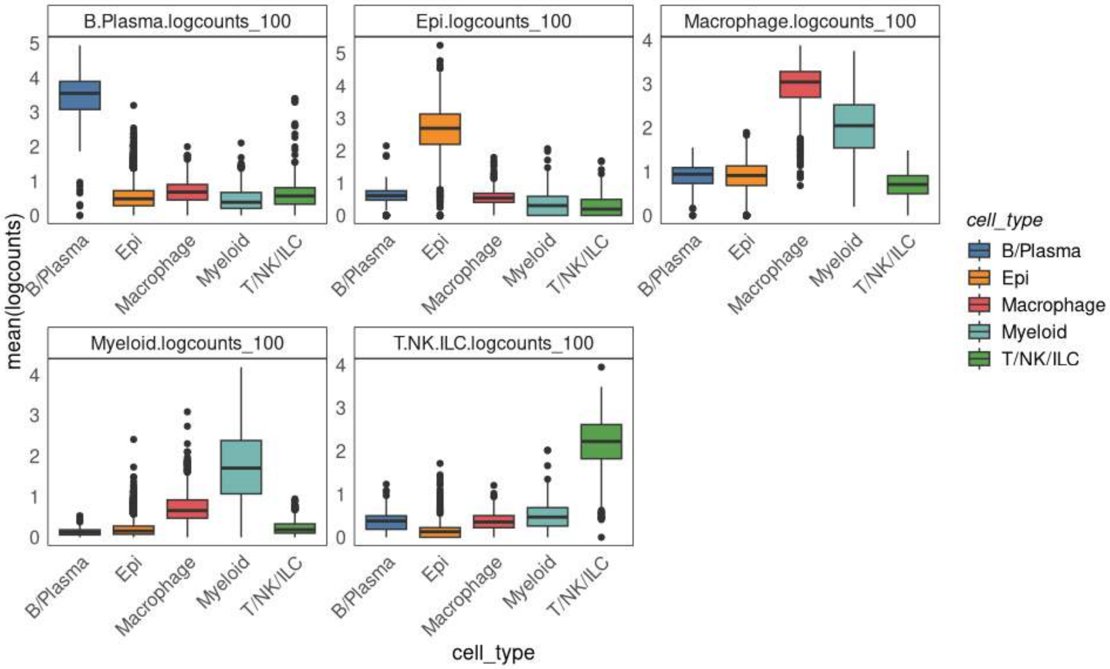


**Supp Figure 7. Average *scran*-normalized logcounts of the top 100 markers per cell type identified by *smartid* on scCOVID19.**

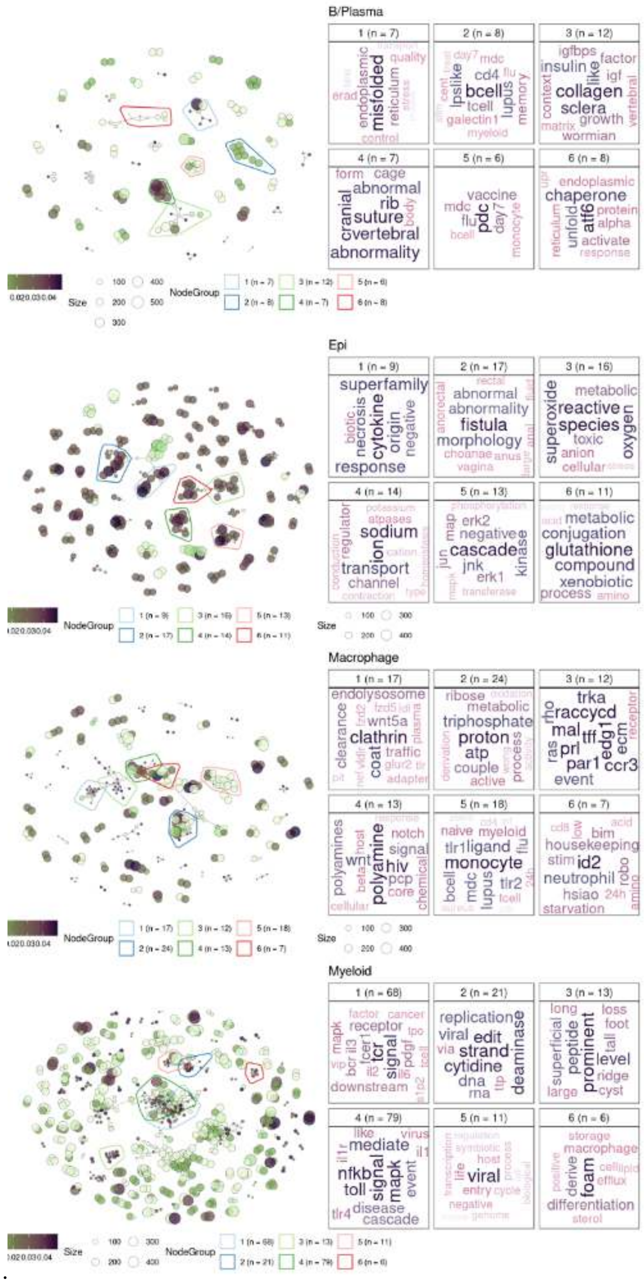


**Supp Figure 8. Network visualization and word clouds of the top 6 clusters for the GSEA results across cell types.**

From top to bottom, the order is B/Plasma, Epithelial, Macrophage and Myeloid cell marker genes.

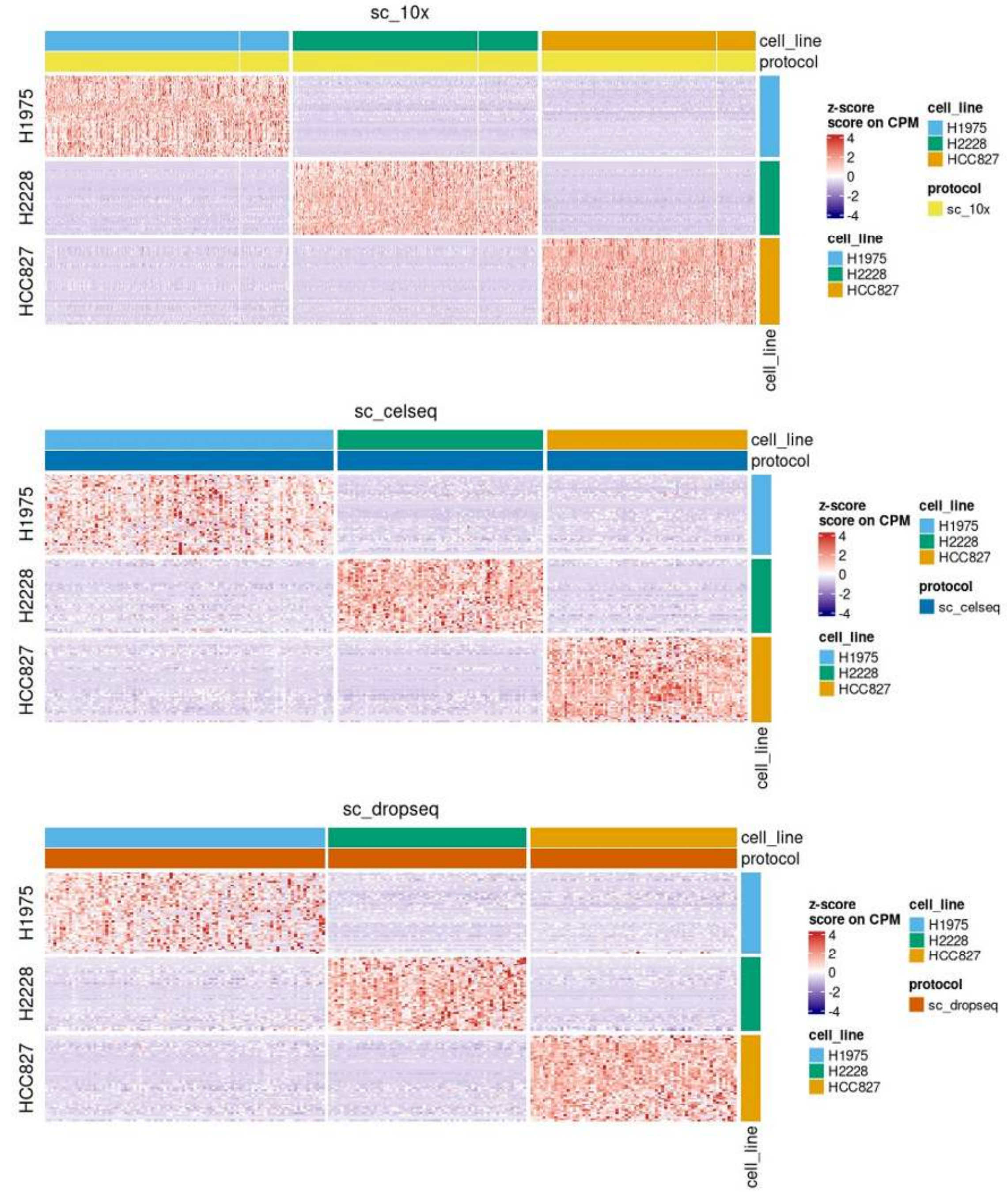


**Supp Figure 9. Heatmap of scaled *smartid* score of the top 50 marker genes on each dataset from platform 10X Chromium (top), CEL-Seq2 (middle) and Drop-Seq (bottom).**

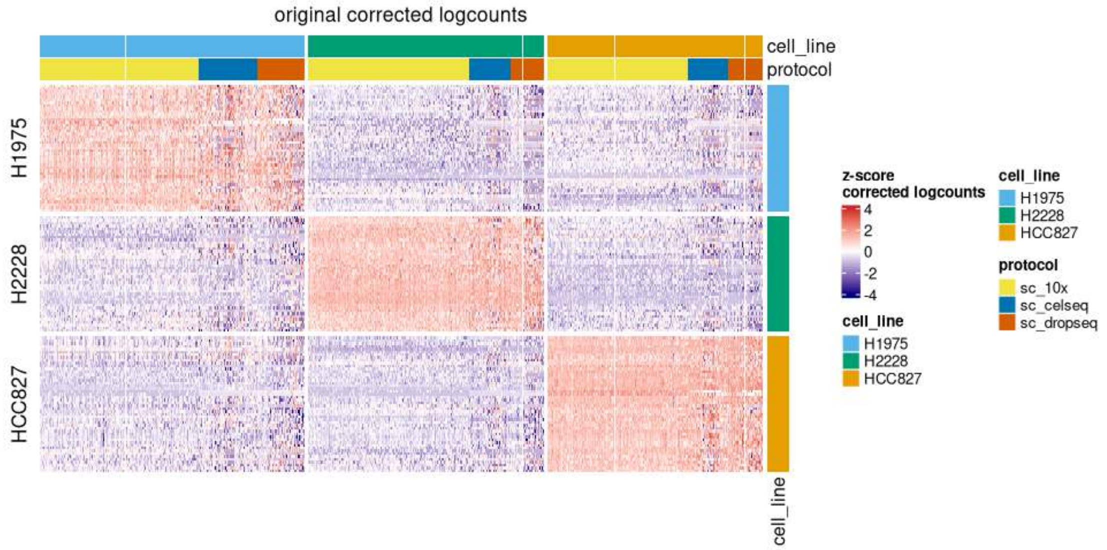


**Supp Figure 10. Heatmap of scaled MNN corrected logcounts of the top 50 marker genes across all cells.**

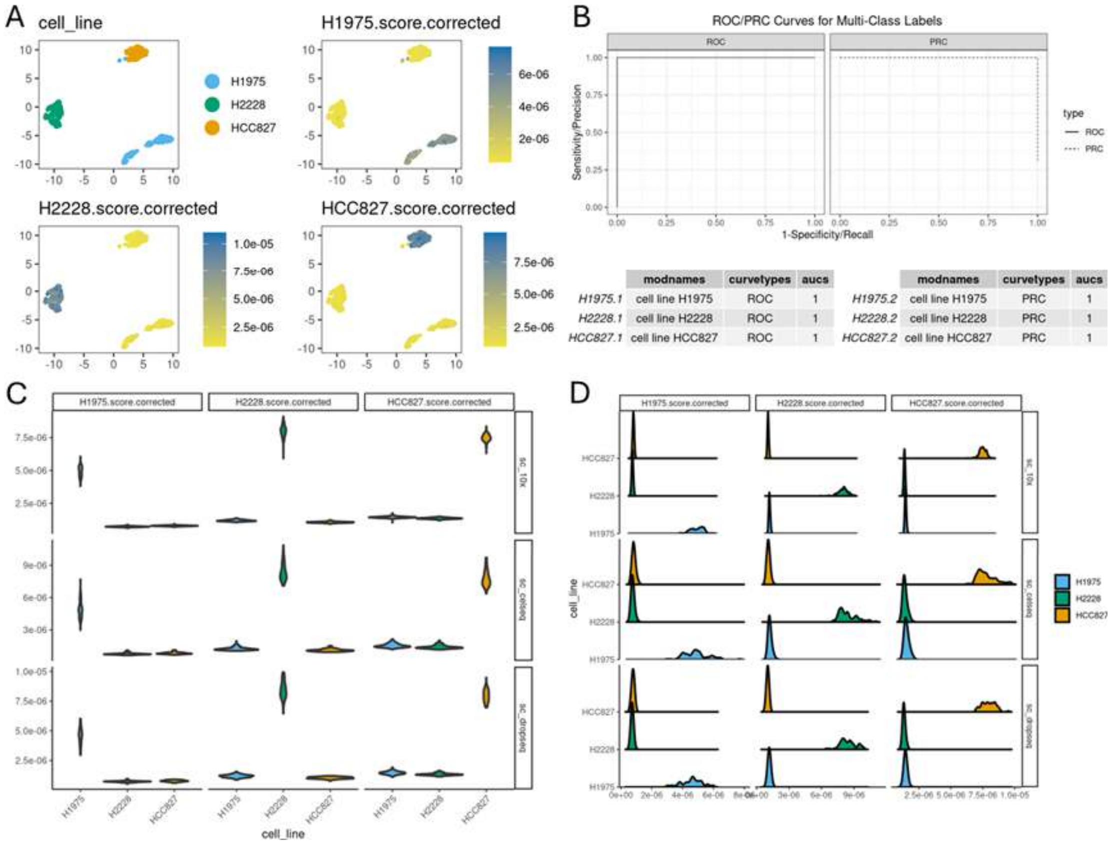


**Supp Figure 11. Clear separation of cell lines by *smartid* gene set score.**

(A) UMAP coloured by cell line (top left) and by *smartid* score of each cell line markers. (B) ROC and PRC with AUC listed below. (C) Violin plot of each cell line marker score, split by cell line score (x) and platform (y). (D) Ridge plot of each cell line score, split by cell line score (x) and platform (y).

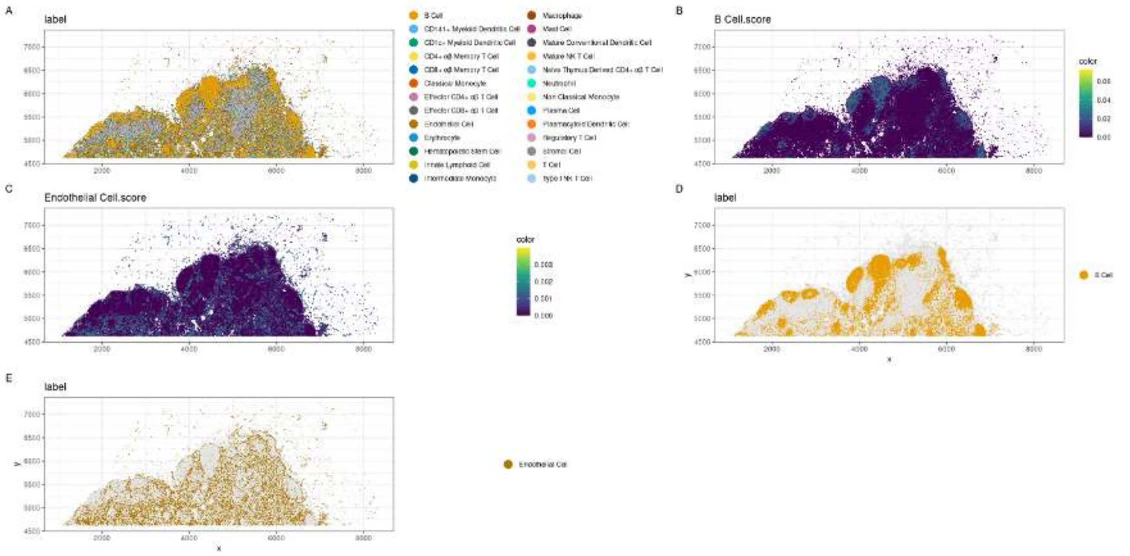


**Supp Figure 12. Cell type label and marker score mapped on spatial slide.**

(A) Spatial map of cell type label. (B) B cell marker score mapped on spatial slide. (C) Endothelial cell marker score mapped on spatial slide. (D) B cell label mapped on spatial slide. (E) Endothelial cell label mapped on spatial slide.

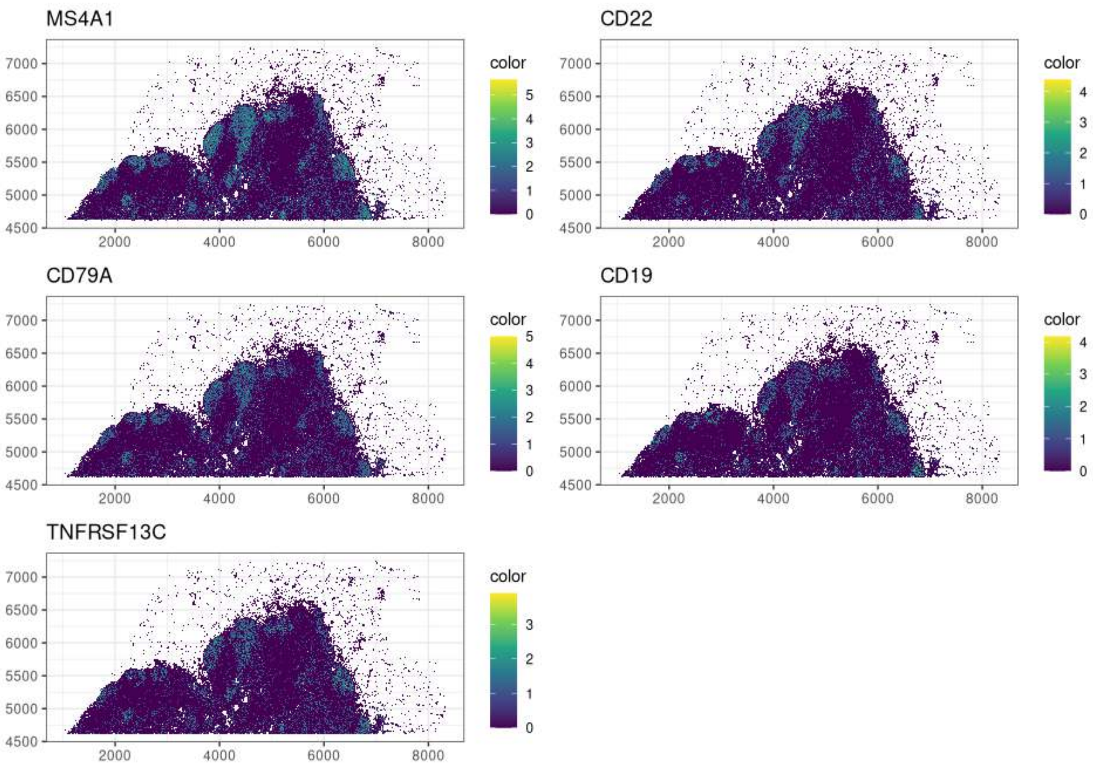


**Supp Figure 13. Spatial distribution of the expression of the five marker genes identified by *smartid* for B cells.**
